# Divergent MEK/ERK and AMPK signaling dictate lipogenic plasticity and dependence on fatty acid synthesis in Glioblastoma

**DOI:** 10.1101/2022.04.07.487530

**Authors:** Katharina M. Eyme, Alessandro Sammarco, Roshani Jha, Hayk Mnatsakanyan, Rudolph Neustadt, Charlotte Moses, Ahmad Alnasser, Daniel Tardiff, Baolong Su, Kevin J Williams, Steven J. Bensinger, Chee Yeun Chung, Christian E. Badr

## Abstract

Deregulated de novo lipid synthesis (DNLS) is a potential druggable vulnerability in Glioblastoma (GBM), a highly lethal and incurable cancer. Yet the molecular mechanisms that determine susceptibility to DNLS-targeted therapies remain unknown, and the lack of brain-penetrant inhibitors of DNLS has prevented their clinical evaluation as GBM therapeutics. Here, we report that YTX-7739, a clinical-stage, brain-penetrant inhibitor of stearoyl CoA desaturase (SCD), triggers lipotoxicity in patient-derived GBM stem-like cells (GSCs) and inhibits fatty acid desaturation in GSCs orthotopically implanted in mice. When administered as a single agent, or particularly in combination with the first line GBM chemotherapy, Temozolomide (TMZ), YTX-7739 showed therapeutic efficacy in orthotopic GSC mouse models owing to its lipotoxicity and its ability to impair DNA damage repair. Leveraging genetic, pharmacological, and physiological manipulation of key signaling nodes in gliomagenesis, we uncover that aberrant MEK/ERK signaling and its repression of the energy sensor AMP-activated protein kinase (AMPK) primarily drives therapeutic vulnerability to SCD and other DNLS inhibitors. Conversely, AMPK activation mitigates lipotoxicity and renders GSCs impervious to the loss of DNLS, both in culture and *in vivo*, by decreasing the saturation state of phospholipids and diverting toxic lipids into lipid droplets. Altogether, our findings reveal mechanisms of metabolic plasticity in GSCs and provide a framework for rational integration of DNLS-targeted therapies for the treatment of GBM.

## Introduction

Glioblastoma (GBM) represents the most frequent, aggressive, and lethal form of gliomas, the most common primary brain tumor in adults ^1^. GBM tumors are enriched with subsets of tumor cells with a cancer stem-like cells (GSCs) phenotype, which significantly contributes to the inherent heterogeneity and therapeutic resistance ^2^. GSCs are particularly endowed with metabolic plasticity and adaptability, which allows them to outcompete mature GBM cells with defined metabolic dependencies, survive in a harsh microenvironmental niche poor in nutrients, and overcome various therapeutic insults ^1^. Unlike the non-stem tumor cells, GSCs can metabolize multiple nutritional substrates and rely on oxidative phosphorylation and de novo lipid synthesis (DNLS) to support their energy demand and self-renewal, thus sustaining tumor growth and proliferation ^1^.

The reprogramming of cellular lipid metabolism is intricately linked with malignant transformation, progression, metastasis, and therapeutic resistance ^2-4^. Oncogenic signaling in GBM, which regulates many such properties and allows GBM cells to cope with their energy demand, is primarily orchestrated by PI3K/AKT and RAS/MAPK signaling ^5,6^. Although RAS mutations are rarely detected in malignant gliomas ^7^, the RAS/RAF/MEK/ERK pathway is aberrantly activated in these tumors ^6^, and ERK phosphorylation is increased in high-grade gliomas ^8^. PI3K/AKT and RAS/MAPK can modulate the activity of various enzymes involved in lipid metabolism, thereby defining the tumor lipidome ^2^, and can regulate critical sensors of cellular metabolic demands such as AMP-activated protein kinase (AMPK), which plays a crucial role in restoring metabolic homeostasis ^9^.

Although various cues and genetic factors can impact the availability of fatty acids (FA) to tumor cells, DNLS is dominantly regulated at the transcriptional level through the activation of sterol regulatory element-binding proteins (SREBPs) ^10^. Stearoyl CoA Desaturase 1 (SCD), a transcriptional target of SREBP, is an enzyme responsible for the conversion of saturated FA (SFA) to monounsaturated FA (MUFA). We have previously shown that GSCs depend on adaptive activation of SCD and that long-term depletion of MUFA in GSCs decreased their self-renewal properties and impaired tumor initiation in mice ^11^. On the other hand, pharmacological inhibition of SCD is primarily toxic due to the accumulation of SFA (a process referred to as lipotoxicity), which induces ER stress and terminal unfolded protein response (UPR) signaling ^11^. Despite its poor ability to penetrate the blood-brain barrier (BBB), the SCD inhibitor CAY10566 showed significant therapeutic efficacy when delivered intranasally ^11^ or systemically ^12^ in mice bearing primary GBM tumors. An orally available SCD inhibitor, YTX-7739, was recently identified as a lead clinical candidate to treat synucleinopathies ^13^. Based on its biochemical potency, safety, favorable pharmacokinetics, and the ability to cross the BBB from rats to cynomolgus monkeys ^13^, YTX-7739 is currently being evaluated in phase I clinical trials for the treatment of patients with Parkinson’s disease (https://www.trialregister.nl/trial/8258; https://www.trialregister.nl/trial/9172). Translational efforts of SCD-targeted therapy for GBM have been hindered by the absence of clinically relevant, brain-penetrant inhibitors of this desaturase. Further, previous work has left unresolved whether precise genetic alterations could drive the dependence of GSCs on SCD or DNLS in general and whether defined molecular markers could predict vulnerability or resistance to DNLS-targeted therapies. Here, we explored the functional and therapeutic efficacy of YTX-7739 in GBM, and we undertook a discovery effort to understand how oncogenic signaling can impact lipid metabolism plasticity in GSCs.

## RESULTS

### A clinical-stage inhibitor of SCD selectively inhibits fatty acid desaturation in patient-derived GSCs in vitro

To test whether YTX-7739 effectively reduces FA desaturation in GSCs by selectively inhibiting SCD, we performed quantitative FA analysis of GSCs after genetic silencing of this desaturase with short-hairpin RNA (shRNA) or treatment with YTX-7739. Unsupervised hierarchical clustering of FA content revealed a comparable profile after drug treatment or SCD knockdown (**Figure 1A**). Consistent with prior studies ^13^, treatment of GSCs with YTX-7739 resulted in a 74% and 81% decrease in the desaturation index (DI) of C16 and C18, respectively (**Figure 1B**). We then measured cell viability after treatment with YTX-7739 in normal human astrocytes (NHA) and 12 patient-derived GSCs lines with distinct genetic signatures isolated from primary and recurrent GBM (**Figure S1A**). NHA were the least sensitive to YTX-7739 with an EC_50_ of 81.41 µM (**Figure 1C**). While sensitivity to YTX-7739 varied among GSCs, 7/12 GSCs lines tested had an EC_50_ < 4µM and were therefore classified as sensitive to this inhibitor, while those with an EC_50_ > 20µM were classified as resistant for subsequent studies (**Figure 1C**). We did not observe any correlation between SCD expression and response to YTX-7739 (**Figure S1B**). Supplementing the cell culture medium with Oleate (C18:1n9), the main product of SCD, but not Palmitate (C16:0; PA), an SCD precursor, rescued from cell death caused by YTX-7739 (**Figure 1D**), thus confirming that the observed cell death is directly caused by SCD inhibition. Further, ectopic expression of SCD in sensitive GSCs reverted YTX-7739 toxicity, further corroborating on-target activity (**Figure 1E**). Overall, YTX-7739 kills GSCs by specifically inhibiting SCD-mediated FA desaturation.

**Figure 1:**
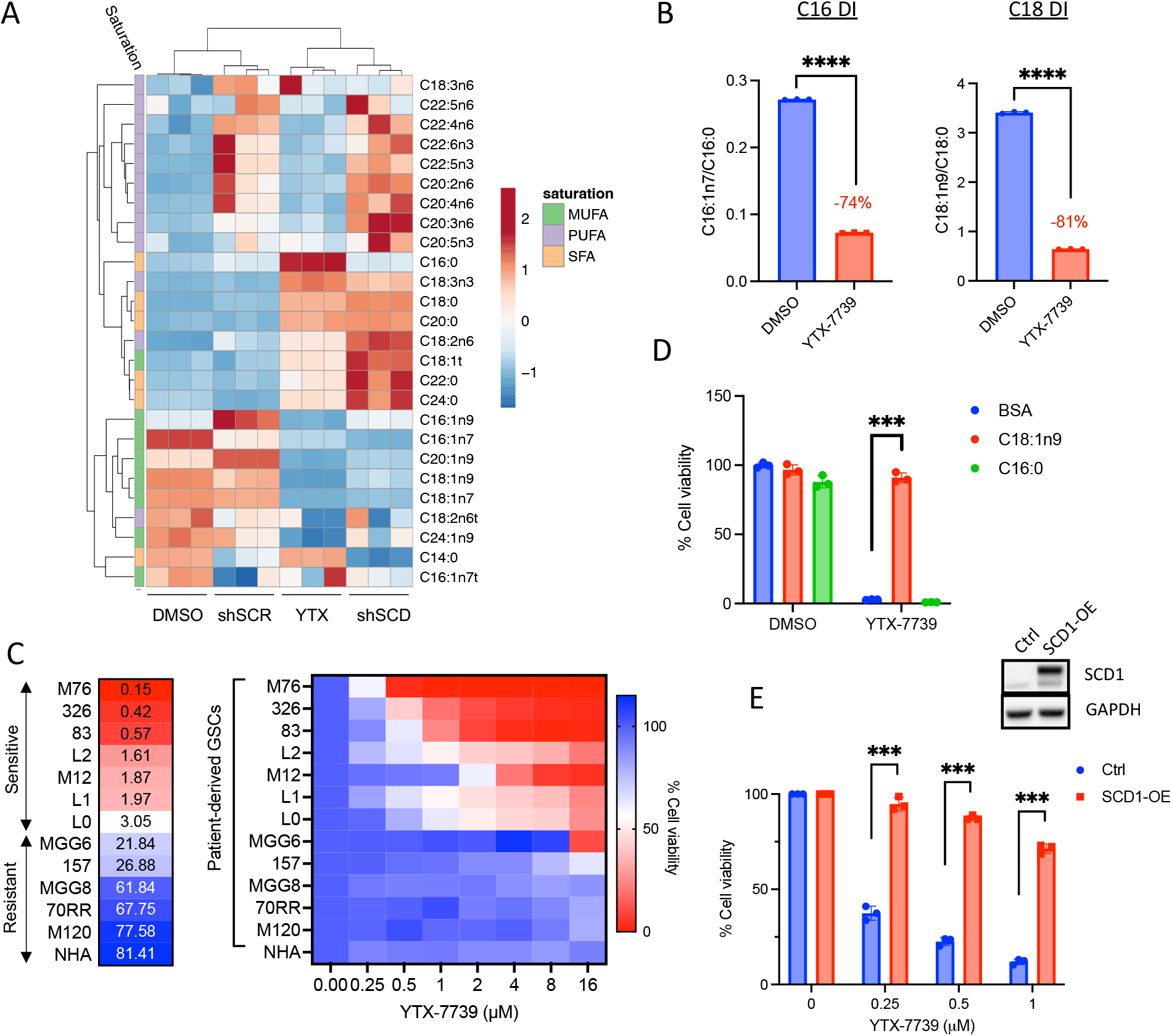
YTX-7739 is a selective inhibitor of FA desaturation in cultured GSCs. (A) Heat map representing unsupervised hierarchical clustering of FA content in 83 GSCs either transduced with a control shRNA (shSCR) or shSCD or treated with DMSO (control) or YTX-7739 (1µM) for 72h. (B) Desaturation index (DI) of C16 and C18 in 83 GSCs treated with YTX-7739 (1µM) for 72h. (C) Cell viability and EC_50_ of NHA and 12 GSCs treated with YTX-7739 at the indicated doses for 96h, expressed as a percentage of control. (D) Cell viability in 83 GSCs treated with YTX-7739 (1µM) in the presence of BSA (control) or the indicated BSA-complexed FA for 96h. (E) Cell viability of 83-GSCs ectopically expressing a control vector (Ctrl) or SCD1 (SCD1-OE) treated with YTX-7739 for 96h. Student t-test ****P<0.0001; *** P<0.001; **P<0.01; *P<0.05

### YTX-7739 selectively kills GBM by exacerbating ER stress and promoting IRE1-JNK mediated cell death

Quantification of individual long-chain SFA revealed that SCD precursors (C16:0 and C18:0) were particularly abundant in GSCs and further enriched after YTX-7739 treatment (**Figure S2A**) suggesting that this inhibitor promotes SFA-mediated lipotoxicity in GSCs. Following treatment with YTX-7739, we observed a marked upregulation of ER stress/UPR markers, including GRP78, spliced XBP1 (sXBP1), C/EBP-homologous protein (CHOP), and its direct transcriptional target GADD34 ^14^, as well as increased phosphorylation of Ser51 on the α-subunit of eukaryotic translation initiation factor 2 (p-EIF2α) (**Figure 2A-B**). We also observed increased phosphorylation of c-Jun N-terminal kinase (JNK), an essential regulator of the UPR-mediated apoptosis ^15,16^, and its downstream target c-Jun (**Figure 2B**). In accord with this, alleviating ER stress with Phenylbutyrate (PBA), Azoramide (AZO), or Salubrinal (SAL) significantly protected against SCD inhibition (**Figure 2C**). Similarly, two inhibitors of the UPR sensor IRE1 and two JNK inhibitors confer protection against YTX-7739 (**Figure 2C and S2B**). Supplementing the cell culture medium with C16:0 to increase endogenous SFA levels strongly impaired cell viability in two GSCs lines (M120 and MGG8) resistant to YTX-7739 (**Figure S2C-D**), which infers that, regardless of the phenotypic outcome, SCD is effectively inhibited in all GSCs. However, lipotoxicity and ensuing cell death only occur when SFA accumulate at cytotoxic levels. If this hypothesis is correct, then concomitant inhibition of SCD along with the upstream enzyme fatty acid synthase (FASN) or acetyl-CoA carboxylase (ACC) (to prevent SFA synthesis; **Figure 2D**) would be rather protective against SCD inhibition. Indeed, the combination of YTX-7739 with a relatively low FASN or ACC inhibitor dose abrogated cytotoxicity (**Figure 2E**). These results confirm that pharmacological inhibition of SCD promotes SFA-mediated lipotoxicity and ER stress activation of IRE1-JNK signaling in GSCs.

**Figure 2:**
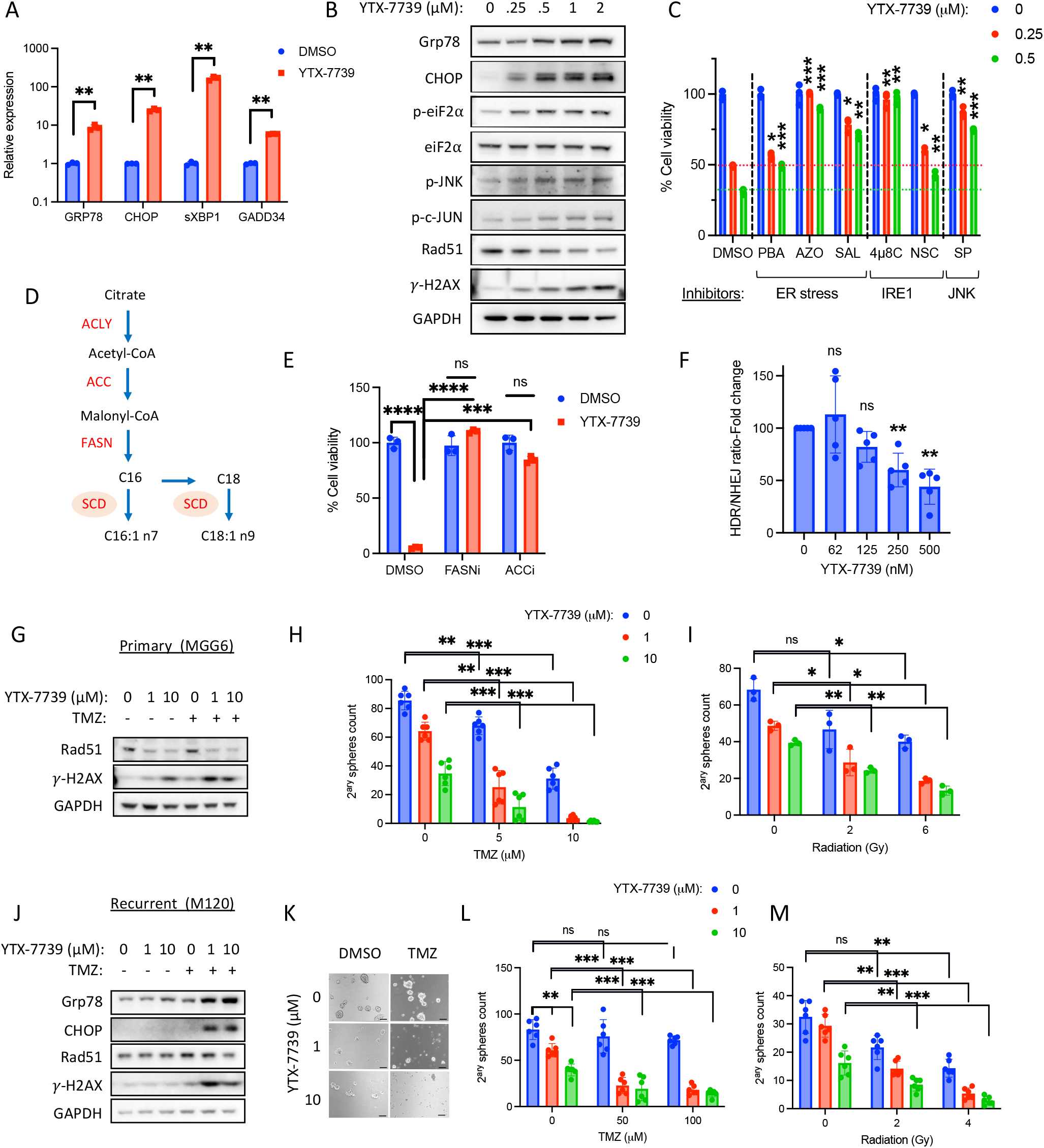
Treatment with YTX-7739 promotes a terminal UPR response and sensitizes to TMZ and radiation. (A) Relative mRNA expression of ER stress markers in 83 GSCs treated with DMSO or YTX-7739 (1µM) for 48h. (B) Immunoblot analysis of 83 GSCs treated with the indicated does of YTX-7739 for 48 h. (C) Cell viability of 83-GSCs treated with YTX-7739 (0.5-1µM) or its combination with inhibitors of ER stress (PBA: 4mM; Azoramide: 50µM; Salubrinal: 25µM), IRE1 (4µ8C: 25µM, NSC95682: 20µM) and JNK (SP600125: 20µM) for 96h. (D) Schematic overview of DNLS pathway. (E) Cell viability of 83 GSCs treated with YTX-7739 (0.5µM) in combination with inhibitors of FASN (GSK2194069; 0.1µM) or ACC (CP-640186; 10µM) for 96h. (F) 83 GSCs stably expressing BLRR system to monitor DNA damage repair were treated with YTX-7739. The BLRR assay is represented as a fold-change in HDR/NHEJ normalized to cell viability. (G-I) MGG6 GSCs (primary) were pretreated with YTX-7739 followed by TMZ (5µM) or RT. (G) Immunoblot analysis of DNA repair and DNA damage at 72h after treatment with YTX-7739 and TMZ (H-I) Secondary spheres were counted on day 9 after co-treatment with TMZ (H) or RT (I). (J-M) M120 GSCs (recurrent) were pretreated with YTX-7739 followed by TMZ (100µM) or RT. (J) Immunoblot analysis after treatment with YTX-7739 and TMZ. (K) Representative micrographs of neurospheres. Scale bar, 100µm. (L-M) Secondary spheres were counted on day 9 after co-treatment with TMZ (H) or RT (I). ****P<0.0001; *** P<0.001; **P<0.01; *P<0.05

### Treatment with YTX-7739 increases sensitivity to radiation and Temozolomide in primary and recurrent GBM in culture

Double-strand breaks (DSB) at DNA targeted sites are typically repaired through error-prone non-homologous end joining (NHEJ) or template-dependent homology-directed repair (HDR)^17,18^ that is mediated by RAD51^19^. In GBM and particularly GSCs, increased RAD51 expression and HDR repair contribute to their inherent and acquired resistance to TMZ^20^ and radiation therapy (RT)^21,22^. We have previously reported that the upregulation of ER stress following pharmacological inhibition of SCD with CAY10566 decreased RAD51 expression, thus sensitizing GSCs to TMZ^11^. Treatment with YTX-7739 also led to a decreased RAD51 expression concomitant with an upregulation of the DNA damage marker γ-H2AX (**Figure 2B**). We used an HDR/NHEJ reporter system ^23^ and confirmed that treatment of GSCs with YTX-7739 significantly decreases the ratio of HDR/NHEJ (**Figure 2F**), in line with RAD51 downregulation. Next, we asked whether sensitization to TMZ or RT could occur in GSCs resistant to the SCD inhibitor or those resistant to TMZ. In these experiments, we assessed cell viability (potentially altered by lipotoxicity or DNA damage) and stem cell properties (measured using a secondary sphere formation assay). Treatment of MGG6 (YTX-7739 resistant; TMZ sensitive) with YTX-7739 alone downregulated RAD51 and increased γ-H2AX, which was further increased when the SCD inhibitor was combined with TMZ (**Figure 2G**). Consequently, such a combination almost entirely abrogated long-term secondary spheres formation and decreased cell viability (**Figures 2H and S2E**). Further, YTX-7739 with RT also significantly decreased secondary spheres count (**Figure 2I**). In the recurrent GSCs line M120 (YTX-7739- and TMZ-resistant), the combination of YTX-7739 with TMZ, but not the SCD inhibitor alone, decreased RAD51 protein expression along with an increase in ER stress (Grp78 and CHOP) and γ-H2AX (**Figure 2J**). A relatively high dose of TMZ (50-100µM) did not significantly alter spheres count or cell viability (**Figure 2K-L and S2F**), and YTX-7739 (at 10 µM) did not change short-term cell viability as expected but significantly decreased sphere count (**Figures 2K-M and S2F**). However, YTX-7739 with TMZ strongly decreased both secondary spheres count and cell viability (**Figures 2K-L and S2F**). Treatment with YTX-7739 also sensitized this recurrent GSC line to RT (**Figure 2M**). Overall, our data raise the prospect of synergy between YTX-7739 and DNA damaging standard-of-care therapy for GBM to increase therapeutic efficacy.

### YTX-7739 inhibits fatty acid desaturation and extends survival in orthotopic GBM mouse models

We assessed the pharmacodynamics (PD) of YTX-7739 in plasma, brain, and tumor tissue in an orthotopic brain tumor mouse model. Animals bearing 83 GSCs brain tumors received a daily intraperitoneal (i.p.) dose of vehicle control or YTX-7739 (30mg/kg) for 11 days before harvesting plasma and brain tissue from both hemispheres to measure the DI in plasma, brain, and brain tumor samples. We observed a significant decrease in C16 and C18 DI in all samples from the YTX-7739 group. Most notably, treatment with YTX-7739 resulted in a 50% and 30% decrease in C16 and C18 DI, respectively, in brain tumors (**Figure 3A**). In conjunction with a decreased DI, we observed a strong upregulation of ER stress markers in brain tumor tissue after treatment with YTX-7739 (**Figure 3B**). To test the therapeutic efficacy of YTX-7739 in GBM mouse models, two GSC lines were selected based on a range of sensitivity to YTX-7739 and TMZ. In a first experiment, mice were implanted with 83-GSCs (highly aggressive; YTX-7739 sensitive - EC_50_ = 0.57 µM, **Figure 1C**; TMZ resistant ^11^) and allowed to establish tumors detected using bioluminescence imaging before initiating treatment. Mice received 16 daily i.p. doses of vehicle (Control group) or YTX-7739 at 10 and 30mg/kg. Albeit modest, treatment with this compound resulted in a significant increase in overall survival (**Figure 3C**). To compare the efficacy of this SCD inhibitor to TMZ, we monitored tumor growth in 83-GSCs-bearing mice treated with vehicle control, YTX-7739 (30mg/kg), TMZ (10mg/kg), or YTX-7739 (30mg/kg) +TMZ (10mg/kg). In line with a modest increase in survival, YTX-7739 as monotherapy did not significantly decrease tumor growth. While TMZ alone initially delayed tumor growth, it was quickly resumed 25 days after tumor implantation (**Figure 3D**). On the other hand, YTX-7739 with TMZ significantly hindered tumor growth compared to TMZ alone (**Figure 3D**). Survival analysis demonstrated that the combination of YTX-7739 with TMZ achieved a significantly longer median survival (43.5 days) than TMZ alone (32 days). Treatment with YTX-7739 or its combination with TMZ was well tolerated throughout the treatment, with no visible signs of discomfort, toxicity, or weight loss (**Figure S3A**). We repeated this experiment with mice implanted with MGG8 GSC (YTX-7739 resistant - EC_50_ = 61.84 µM, **Figure 1C**; TMZ sensitive). A significant decrease in C16 and C18 DI from the plasma of mice treated with YTX-7739 confirmed its effect on circulating fatty acid levels (**Figure S3B**). Longitudinal imaging of tumor growth showed a divergent response in the YTX-7739-treated group, whereby 4/8 mice showed a relatively stable tumor signal up to 10 weeks after tumor implantation, indicative of a strong therapeutic response (**Figure 3F**). In comparison, all mice in the control group (8/8) showed an increased tumor growth over time, and none were alive by week 10. This translated into a significantly extended survival of 91 days in the YTX-7739 treated group compared with 52 days in the control group (**Figure 3G**).

**Figure 3:**
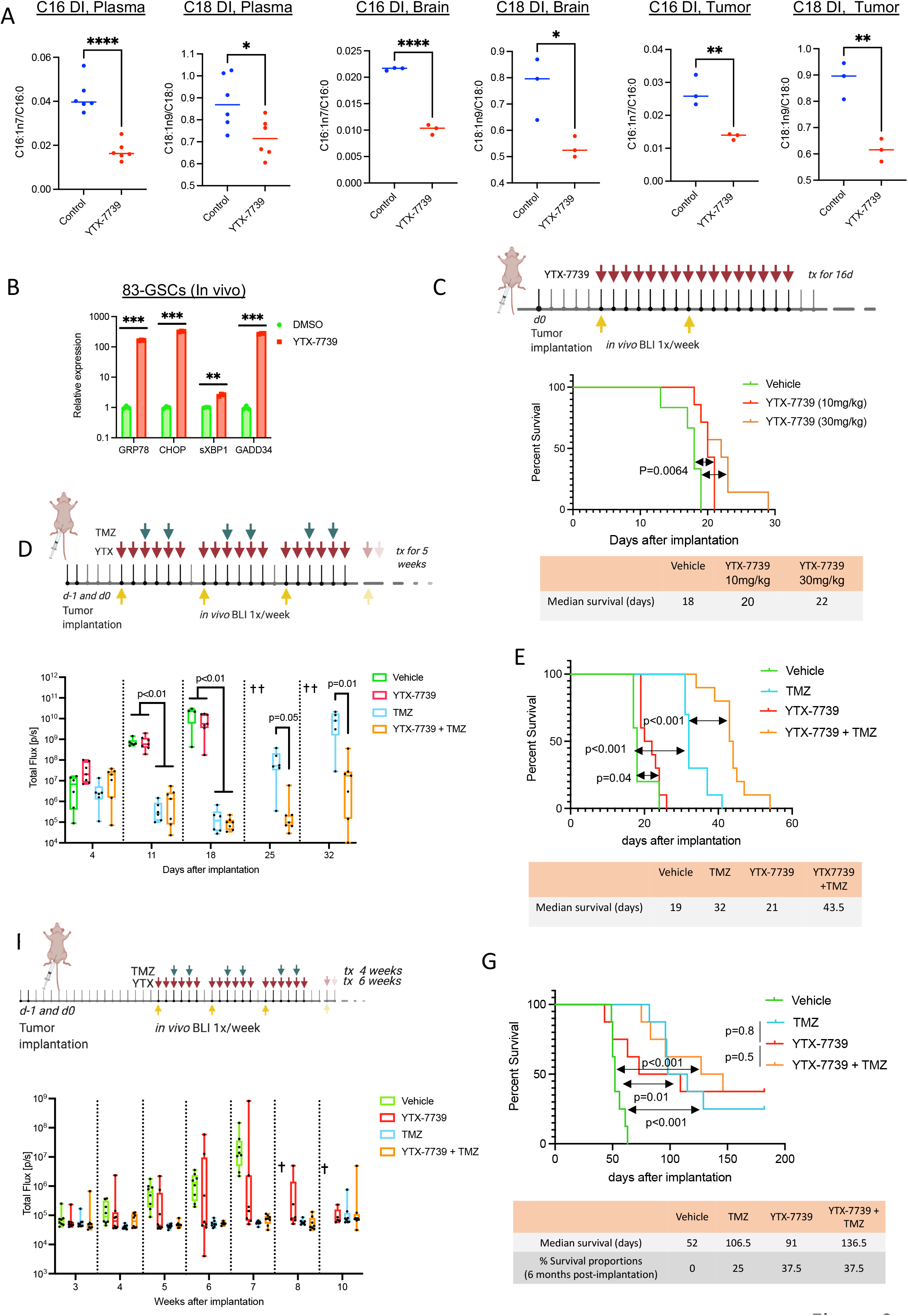
YTX-7739 inhibits fatty acid desaturation and delays tumor growth in patient-derived orthotopic GSCs Models. (A) DI of C16 and C18 in plasma, brain, and brain tumor samples from mice bearing 83-GSCs tumors and treated with YTX-7739 (30mg/kg) for 11 days (n=3 mice/group). (B) Brain tumor tissue from the same experiment (n=1 mouse/group) was also used to determine the relative mRNA expression of ER stress markers. (C) Mice implanted with 83-Fluc GSCs (n = 6/group) received a daily i.p. dose of vehicle (Control) or YTX-7739 (10-30 mg/kg) for 16 days starting at day 5 post-implantation. Kaplan-Meier curves show median survival in all groups (*p = 0.0064, two-sided log-rank test). (D) Overtime monitoring of tumor growth with Fluc imaging in mice implanted with 83-Fluc GSCs and treated with a daily i.p. dose of vehicle (Control; n=6), YTX-7739 (30mg/kg; n=7), TMZ (25mg/kg; n=6) of YTX-7739+TMZ (n=7)). Cross indicates that at least 50% of mice have expired due to tumor burden. (E) Kaplan-Meier curves showing median survival in mice bearing 83-Fluc GSCs and treated with vehicle (Control; n=5), YTX-7739 (30mg/kg; n=10), TMZ (10mg/kg; n=10 or YTX-7739+TMZ (n=10). (F-G) Mice implanted with MGG8-Fluc GSCs (n = 8/group) received a daily i.p. dose of vehicle (Control), YTX-7739 (30 mg/kg), TMZ (5mg/kg) or their combination as indicated. (F) Overtime monitoring of tumor growth with Fluc imaging. (G) Kaplan-Meier curve shows median survival in all groups (p, two-sided log-rank test). ****P<0.0001; *** P<0.001; **P<0.01; *P<0.05

Additionally, the combination of TMZ with YTX-7739 extended survival for 136.5 days compared to 106.5 days for TMZ alone. Despite these GSCs being highly sensitive to TMZ, there was no significant difference in overall survival between mice treated with either TMZ or YTX-7739 alone, suggesting a comparable therapeutic efficacy between YTX-7739 and the standard of care chemotherapeutic. Further, the survival percentage at six months after tumor implantation was 37.5% in mice treated with YTX-7739 alone compared to 25% in the TMZ-treated group. These mice did not show any detectable tumors by bioluminescence imaging at 6 months when this experiment was terminated (**Figure S3C**). Finally, long-term monitoring for up to 6 months did not reveal any toxicity or weight loss in any treated animals (**Figure S3D**). These data demonstrate that YTX-7739 can cross the BBB, functionally inhibit SCD-mediated desaturation in brain tumors, and exert therapeutic efficacy in preclinical mouse models of GBM.

### Signaling through RAS/MEK/ERK promotes vulnerability to SCD inhibitors

Given the disparate sensitivity to SCD inhibitors between NHA and GSCs (**Figure 1C**) and to define the precise genetic alterations driving vulnerability to SCD inhibitors, we used an astrocyte-based model of gliomagenesis initially described by Sonoda et al. ^24^. In this model, NHA are first engineered to express hTERT, combined with the loss of functional p53 and pRb pathways, to prevent senescence and increase cell proliferation. These immortalized non-transformed astrocytes are hereafter referred to as HA-NT. Ectopic expression of a mutant, constitutively active HRAS^G12V^ (HA-Ras) but not a truncated active form of EGFR (HA-EGFRvIII) in HA-NT promotes anchorage-independent growth (**Figure S4A**) and the ability to form tumors akin to a grade III anaplastic astrocytoma following intracranial implantation in mice ^24^. We confirmed increased phosphorylation of ERK1/2 and AKT in HA-Ras cells and the activation of the truncated EGFR and AKT phosphorylation in HA-EGFRvIII (**Figure 4A**). Next, we examined sensitivity to two SCD inhibitors (YTX-7739 and CAY10566) in NHA and this isogenic astrocyte model. Pharmacological inhibition of SCD led to a minor decrease in cell viability in NHA and HA-NT (**Figure 4B**). Although EGFR signaling has been reported to sensitize GBM cell lines to inhibitors of fatty acid synthesis^9^, HA-EGFRvIII showed a similar response to HA-NT. In contrast, HA-Ras cells were particularly sensitive to SCD inhibitors with an EC_50_ of 209 nM and 18.6 nM for YTX-7739 and CAY10566, respectively (**Figure 4B-C**). This selective toxicity towards HA-Ras was further confirmed through increased apoptosis detected by caspase activation and Annexin V staining (**Figure S4B-C**). ER stress/UPR, DNA damage, and JNK activation were only observed in HA-Ras but not in HA-NT following SCD inhibition (**Figure 4D-E and S4D**). Cell death caused by SCD inhibitors could be reversed by supplementing HA-Ras with Oleate or inhibiting ER stress, IRE1, or JNK, similarly to what is observed in GSCs (**Figure S4E-F**). This implies that the molecular underpinnings of vulnerability and cell death triggered by SCD inhibition are similar in HA-Ras and GSCs. Unexpectedly, we also found that ectopic expression of HRAS, but not EGFRvIII, also increased sensitivity to FASN and ACC inhibitors (**Figure S4G**), thus implying that RAS activation creates an overall dependence on DNLS. Since RAS primarily signals through MEK/ERK, blocking this pathway in HA-Ras with two specific MEK and ERK inhibitors (AZD8330 and AZD0364, respectively) prevented cell death caused by SCD inhibitors (**Figure S4F**), thus reverting this acquired sensitivity. Importantly, we found that inhibiting MEK/ERK in GSCs prevented ER stress and terminal UPR (**Figure 4F-G**) and the phosphorylation of IRE1, JNK, and c-Jun (**Figure 4G**). Consequently, inhibiting MEK/ERK averts PARP cleavage, an indicator of apoptosis, and protects against SCD inhibitors-induced cell death in three GSCs lines (**Figure 4G-H and Figure S4H)**. Finally, ectopic expression of HRAS^G12V^ in one of the YTX-7739 resistant GSCs lines (MGG6) promotes vulnerability to this SCD inhibitor with a 40% increase in cell death after YTX-7739 treatment (**Figure S4I**). Altogether, these results indicate that acquired or inherent sensitivity to SCD inhibitors is primarily mediated by MEK/ERK activity.

**Figure 4:**
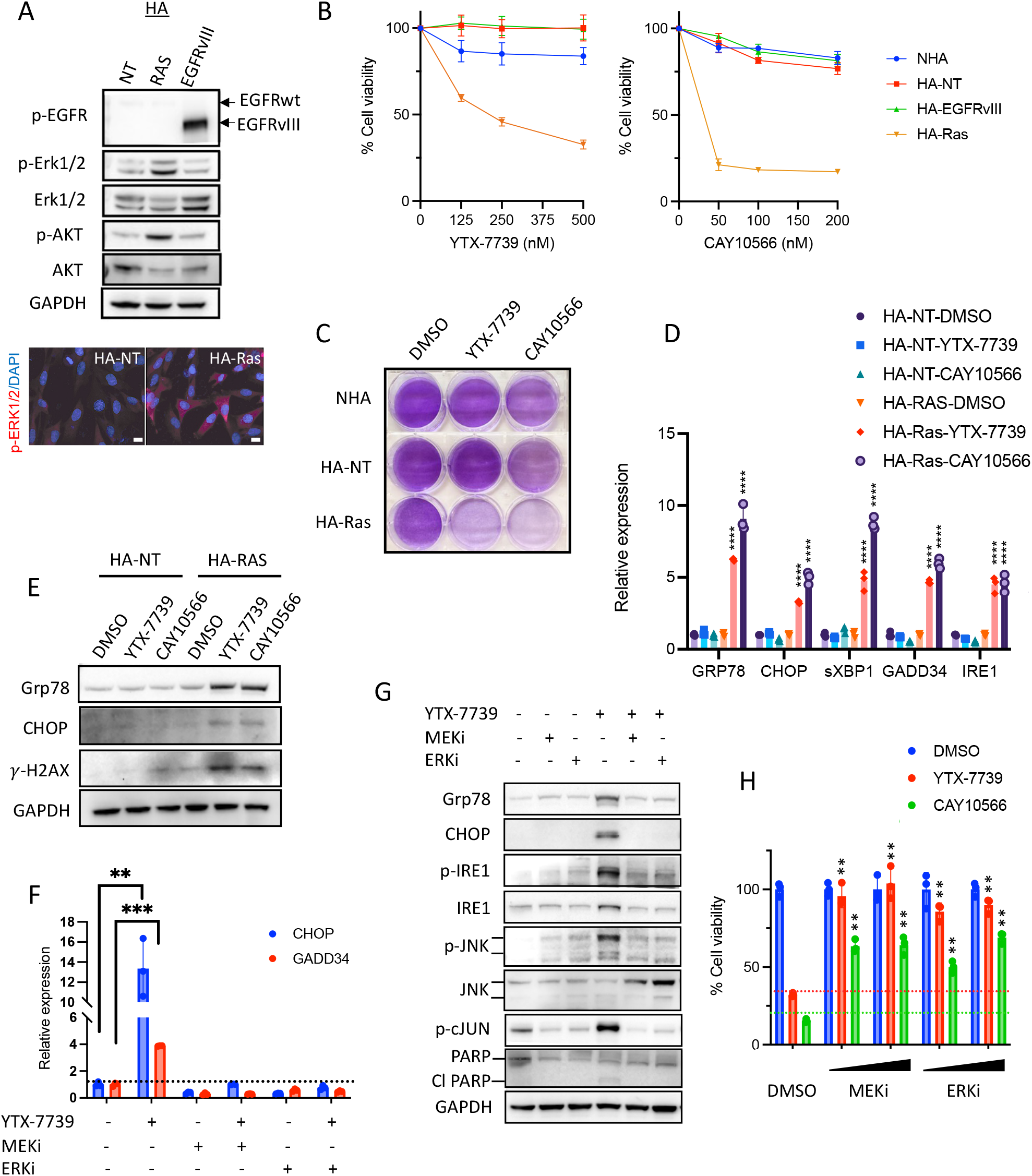
Activation of RAS/MEK/ERK promotes vulnerability to SCD inhibitors. (A) Immunoblot analysis and immunostaining of HA-NT cells stably expressing a control vector, HRAS^G12V^ or EGFRvIII. Scale bar, 10µm. (B) Cell viability of primary (NHA) and transformed astrocytes (in 1% serum) treated with YTX-7739 or CAY10566 for 96h. (C) Cells were plated (in 1% serum) and treated with YTX-7739 (1µM) or CAY10566 (0.5µM) for 5 days before staining with crystal violet. (D-E) Relative mRNA expression (D) and immunoblot analysis (E) of ER stress, UPR, and DNA damage in HA-NT and HA-RAS treated with YTX-7739 (1µM) or CAY10566 (0.5µM) for 48 h. (F-G) Relative mRNA expression (F) and immunoblot analysis (G) of ER stress, UPR, JNK, and apoptosis in 83 GSCs treated with YTX-7739 (1µM) or its combination with AZD8330 (MEKi, 1µM), AZD0364 (ERKi, 1µM) for 72 hours. (H) Cell viability of 83 GSCs treated with YTX-7739 (1µM), CAY10566 (0.5 µM), or their combination with MEKi and ERKi (0.25-0.5µM) for 96 hours. ****P<0.0001; *** P<0.001; **P<0.01; *P<0.05

### GSC subpopulations with low MEK-ERK signaling can escape DNLS-targeted therapy

Given the molecular heterogeneity and genetic plasticity of GSCs, we reasoned that cell subpopulations with low MEK-ERK activity could escape SCD-targeted therapy. To explore this possibility, we subjected two GSC lines that are sensitive to (83 and M76) to several rounds of treatment with this inhibitor (YTX-7739 was added twice weekly at 1μm for 6 weeks). Whereas most cells cultured under such conditions expectedly failed to survive SCD inhibition, few cells, referred to as 83R and M76R, could be propagated over an extended period. Rechallenging these cells with either YTX-7739 or CAY10566 resulted in lower cell death (**Figure 5A and S5A**) and an attenuated UPR (**Figure 5B and S5B**) as compared to their parental counterparts. In 83R cells, and to a lesser extent in M76R, we also observed a decreased sensitivity to FASN and ACC inhibitors (**Figure S5C**). Interestingly, along with this increased resistance to DNLS inhibitors, we observed reduced phosphorylation of ERK1/2 and its downstream target RSK1 in resistant cells compared to their parental lines (**Figure 5C**). Finally, ectopic expression of HRAS^G12V^ in 83R restored sensitivity to YTX-7739 (**Figure S5D**). These results support an essential role of MEK-ERK signaling in defining sensitivity and resistance to SCD and DNLS inhibitors.

**Figure 5:**
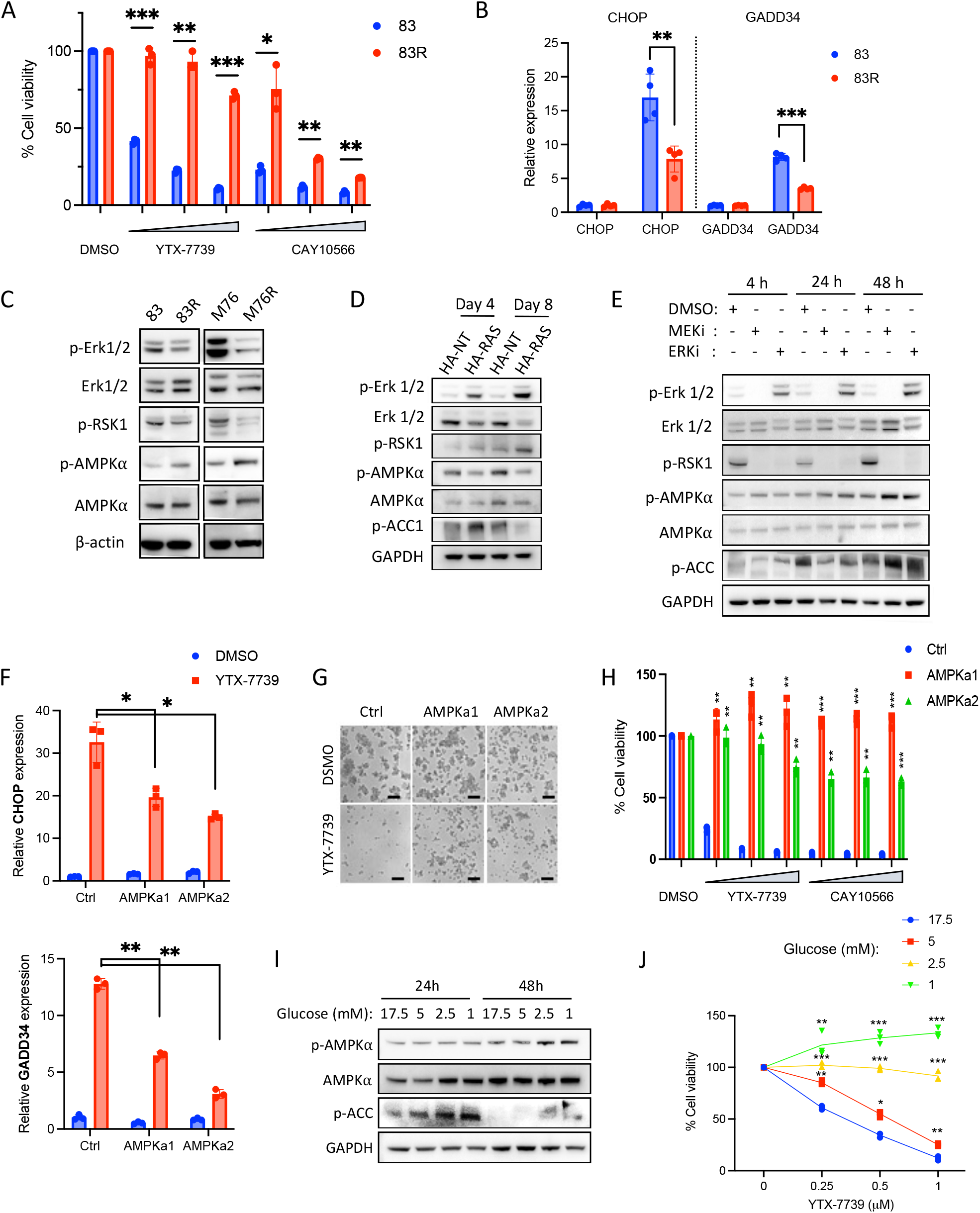
MEK-ERK signaling inhibits AMPK activity and defines the therapeutic response to DNLS inhibitors. (A) Cell viability of 83 and 83R treated with YTX-7739 (0.25-1µM) and CAY10566 (0.125-0.5µM) for 96h. (B) Relative mRNA expression of UPR markers CHOP and GADD34 in 83 and 83R GSCs treated with DMSO (control) or YTX-7739 (1µM) for 72h. (C) Immunoblot analysis of ERK and AMPK signaling in 83/83R and M76/M76R GSCs. (D) Immunoblot analysis of ERK and AMPK signaling in transformed astrocytes on days 4 and 8 after expression of a control (HA-NT) or HRAS^G12V^ (HA-RAS). (E) Immunoblot analysis of ERK and AMPK activity in 83 GSCs treated with MEK and ERK inhibitors. (F) Relative mRNA expression of UPR markers CHOP and GADD34 in 83 Ctrl, AMPKa1, and AMPKa2 treated with DMSO (control) or YTX-7739 (1µM) for 72h. (G) Representative micrographs of 83 GSCs expressing Ctrl, AMPKa1, or AMPKa2 after treatment with YTX-7739 (1µM) Scale bar: 100µm. (H) Cell viability of M76 GSCs after treatment with YTX-7739 (0.25-1µM) or CAY10566 (0.125-0.5µM) for 96h. (I) Immunoblot analysis of AMPK signaling in 83 GSCs cultured in decreasing concentration of Glucose for 24-48h. (J) Cell viability of 83 GSCs cultured in decreasing concentration of Glucose and concomitantly treated with YTX-7739 for 72h. (K) immunoblot analysis of ERK and AMPK activities in MGG6 after supplementing with insulin for 1h. (L) Cell viability of MGG6 and MGG8 treated with YTX (1µM) in the presence of an increasing concentration of insulin for 96h.

### AMPK signaling is repressed by MEK-ERK activation

Our effort to identify regulators downstream of MEK-ERK prompted us to consider AMPK, given its previously reported roles in phosphorylating and inactivating ACC ^25^, promoting a catabolic switch ^26^, and suppressing JNK activity ^27,28^. We observed increased AMPK phosphorylation in 83R and M76R cells compared to their parental cells (**Figure 5C**). AMPK signaling is reportedly downregulated after constitutive activation of Ras, Raf, or MEK/ERK pathway ^29-31^. This was confirmed in our astrocytes model, where we saw a decreased phosphorylation of AMPK and ACC, notably at day 8 post-transduction with HRAS^G12V^-expressing lentivirus (**Figure 5D**). Conversely, in 83 GSCs treated with MEK or ERK inhibitors, we saw time-dependent phosphorylation of AMPK and ACC (**Figure 5E**). Of note, treatment with the ERK inhibitor AZD0364 unexpectedly increased ERK phosphorylation, as previously reported for several ERK inhibitors and attributed to feedback activation ^32-34^. However, a complete blockade of MEK/ERK signaling is reflected through a potent downregulation of RSK1 phosphorylation (**Figure 5E**). Overall, these results confirm that MEK/ERK signaling inhibits AMPK activity.

### Therapeutic response to DNLS inhibitors is dictated by AMPK

To test whether vulnerability to DNLS inhibitors is directly mediated by AMPK, we explored genetic, pharmacological, and physiological modulation of AMPK activity. First, we expressed two constitutively active mutants of AMPK (AMPKa1 ^35^ and AMPKa2 ^36^) in two YTX-7739 sensitive GSCs and HA-Ras. AMPK activation significantly dampened or suppressed terminal UPR induced by YTX-7739 (**Figure 5F and S6A-B**) and rendered both GSCs and HA-Ras cells almost entirely insensitive to SCD, FASN, and ACC inhibitors (**Figures 5G-H and S6C-D**). Similarly, pharmacological activation of AMPK with the synthetic ligand A-769662 provided a dose-dependent protection against YTX-7739 in GSCs and HA-Ras cells (**Figure S6E-F**). A second AMPK activator (AICAR) also conferred protection from the SCD inhibitor, while an AMPK inhibitor (dorsomorphin) increased the cytotoxicity of YTX-7739 in two GSC lines (**Figure S6G**). Further, treatment of 83R with dorsomorphin restored sensitivity to YTX-7739 (**Figure S6H**). Dorsomorphin is a non-specific inhibitor of AMPK ^37^, prompting us to explore shRNA-mediated silencing of AMPKA1 and AMPKA2 as an alternative. However, in line with the essential role of AMPK activity in GSCs^38^, cell viability was rapidly and drastically decreased following AMPK silencing (**Figure S6I**), which precluded further testing. Since AMPK has been reported to be activated by glucose deprivation ^39^, we also tested whether activating endogenous AMPK by lowering glucose concentrations in the GSC culture medium (which contains 17.5mM of Glucose) would also protect against SCD inhibition. Notwithstanding a relatively modest increase in AMPK and ACC phosphorylation at the lowest glucose concentrations (**Figure 5I**), we observed gradual and potent protection from SCD inhibitors in 83 and M76 cells (**Figure 5J and Figure S6J**). Insulin promotes sustained activation of AKT and ERK in primary GBM cultures ^40^ (**Figure S7A**) and reduces AMPK activity ^41^. By increasing insulin levels in the growth culture medium (1-10 µg/ml; B27 supplement used in GSC medium already contains insulin), we were able to sensitize two resistant GSCs (MGG6 and MGG8) as well as 83R to YTX-7739, with 20-65% increase in cell death (**Figure S7B-C**). The effect of insulin was reversed following ectopic expression of AMPK (**Figure S7D**), thus confirming that a decreased AMPK activity directly causes insulin-mediated sensitization. Our results show that AMPK activation drives resistance to SCD and DNLS inhibitors while targeting AMPK increases the therapeutic response.

### GSCs adapt to lipotoxicity and impaired DNLS by activating AMPK

We reasoned that AMPK activation could protect against SFA-triggered lipotoxicity. Indeed, constitutive activation of AMPK strongly repressed terminal UPR caused by high levels of C16:0 (**Figure S8A**) and almost fully protected against C16:0 lipotoxicity in two GSC lines (**Figure S8B**). Despite the reported (inactivating) phosphorylation of ACC by AMPK, there was no significant decrease in cell proliferation or the sphere-forming ability of GSCs expressing AMPKa1 or AMPKa2. Further, AMPK activation was unexpectedly protective against FASN and ACC inhibition **(Figure S6D)**, which, unlike SCD inhibition, does not promote lipotoxicity. This suggested that, in addition to being protected against lipotoxicity, GSCs with activated AMPK could become refractory DNLS. Indeed, long-term silencing of SCD (30 days) completely impaired sphere formation in the control group but only partially in AMPKa1 or AMPKa2 expressing GSCs (**Figure 6A**). Silencing of SCD also wholly prevented tumor growth in 83 GSCs following intracranial implantation in mice ^11^, while AMPKa1 expression did not significantly alter tumor growth (**Figure 6B**). Strikingly, and despite a delayed tumor growth, mice in the AMPKa1/shSCD group developed tumors that gradually grew over time (**Figure 6C**).

**Figure 6:**
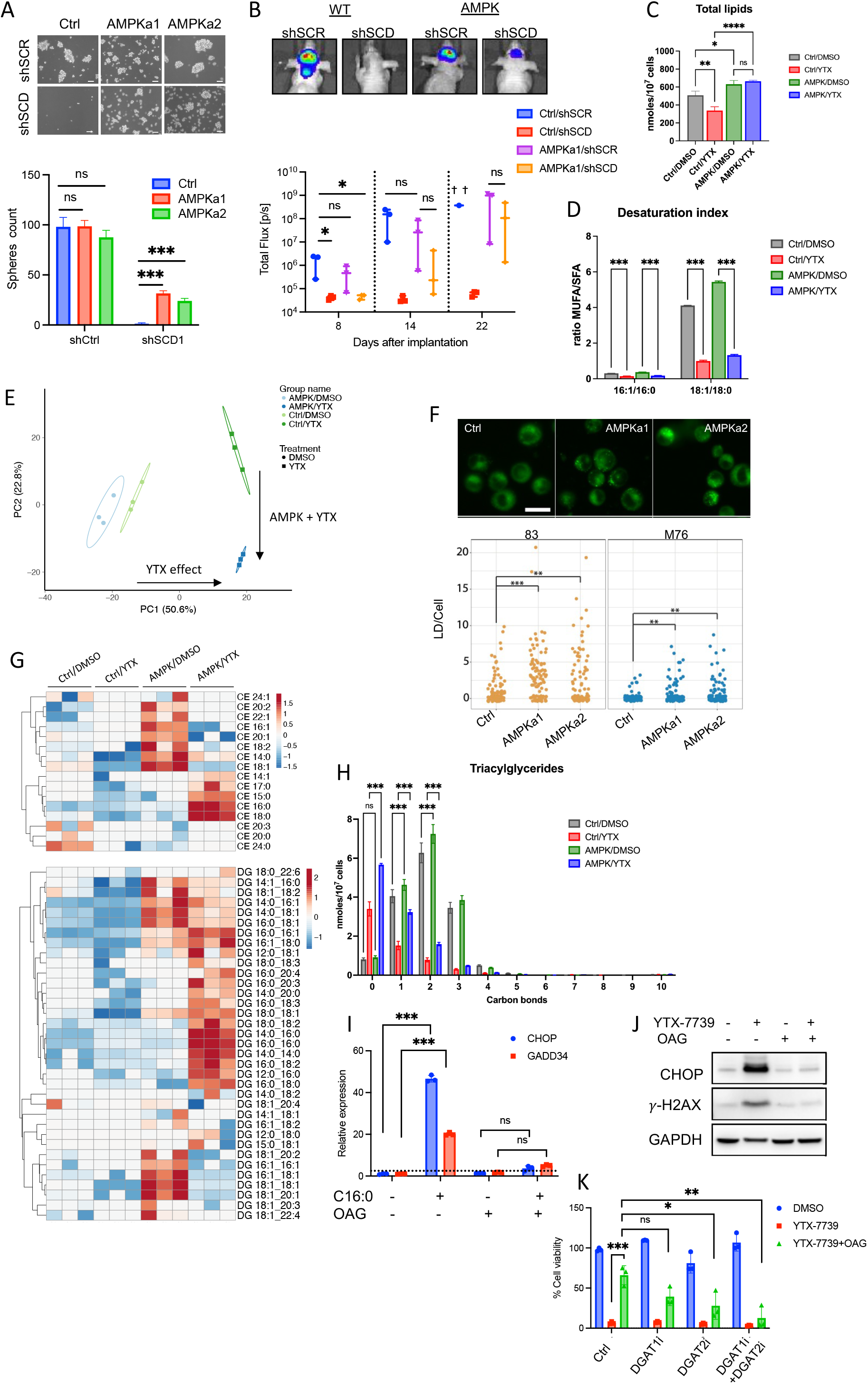
AMPK signaling protects against SCD inhibition and lipotoxicity by altering lipid composition. (A) Representative micrographs and spheres count in 83 GSCs Ctrl, AMPKa1, and AMPKa2 30 days following two rounds of transduction with shCtrl or shSCD lentivirus. Scale bar, 50µm. (B-C) 83-Fluc Ctrl or AMPKa1 GSCs were transduced with shCtrl or shSCD1 and intracranially implanted in mice (n=3) after 48h. (B) Representative images and longitudinal Fluc imaging of mice in each group. Survival analysis is shown using Kaplan-Meier curves. p = 0.0025 (two-sided log-rank test). (C) Barplot representing the amount, expressed in nanomoles/10^7^ cells, of total lipids in untreated (DMSO) and treated (YTX) GSCs Ctrl/AMPK measured by shotgun lipidomics. (D) Barplot representing desaturation indexes of 16:0 and 18:0 in untreated (DMSO) and treated (YTX) GSCs Ctrl/AMPK. (E) Principal component analysis (PCA) of individual lipid species quantified by shotgun lipidomics from untreated (DMSO) and treated (YTX) GSCs Ctrl/AMPK. Percentage of total variance explained by individual principal components (PC1 and PC2). (F) Representative fluorescence imaging of LDs stained with BODIPY 493/503. Scale bar, 25 µm. LDs were quantified in 100 cells of 83 Ctrl, AMPKa1, and AMPKa2. (G) Heatmap of cholesterol esters (CE) and diacylglycerols (DG) quantified by shotgun” lipidomics analysis of untreated (DMSO) and treated (YTX) GSCs Ctrl/AMPK. (H) Barplot representing the amount, expressed in nanomoles/10^7^ cells, of triacylglycerides (TG) with 0-10 carbon bonds in untreated (DMSO) and treated (YTX) GSCs Ctrl/AMPK. (I) Relative mRNA expression of UPR markers CHOP and GADD34 in 83 GSCs supplemented with BSA or C16:0 (300µM) for 24h with or without OAG (60µM). (J) Immunoblot analysis of 83 cells treated with YTX-7739 (1µM) with or without OAG. (K) Cell viability of M76 co-treated with YTX-7739 (0.5µM) and OAG (60µM) in addition to DGAT1i (A922500; 5µM) and/or DGAT2i (PF-06424439; 50 µM) for 96h. *** P<0.001; **P<0.01; *P<0.05

### AMPK signaling protects from SCD inhibition by altering lipid composition

To understand how AMPK mitigates lipotoxicity and impacts GSC lipogenesis, we performed shotgun lipidomics on GSCs expressing a control vector (Ctrl) or AMPKa1 (AMPK) and treated with vehicle (DMSO) or YTX-7739 for 72 hrs. We quantified approximately 900 lipid species across 17 lipid subclasses. We found that enforced AMPK signaling modestly increased the total amount of lipids per cell (**Figure 6C and Figure S8C)**. Treatment of Ctrl GSCs with YTX-7739 significantly decreased the total amount of lipids per cell. Notably, AMPK signaling attenuated the YTX-7739-mediated decrease in lipid amounts (**Figure 6C**).

As expected, examination of acyl tail composition in the lipidome confirmed that YTX-7739 treatment significantly decreased MUFA incorporation into complex lipids and the desaturation index (i.e., 16:1/16:0 and 18:1/18:0), irrespective of AMPK status (**Figure 6D**). Thus, these data show that SCD inhibition alters both lipid desaturation state and total lipid accumulation in GSCs. Next, we applied Principal Component Analysis (PCA) to the lipidomics data to better understand the lipids most contributing to the observed variance between the treatments. PCA revealed minimal differences between untreated Ctrl and AMPK GSCs (**Figure 6E**). Principal component 1 (PC1) captured approximately 50% of the total variance in the lipidomics data and primarily reflected changes to the lipidome in response to YTX-7739 treatment of cells. Principal component 2 (PC2) delineated approximately 23% of the total variance and reflected changes to the lipidome induced by SCD inhibition combined with AMPK signaling. As expected, the top 50 lipids positively contributing to PC1 were enriched for phospholipids containing two saturated acyl tails (e.g., PG 16:0_18:0, PC 16:0_18:0, PE 16:0_16:0) and phospholipids containing one saturated acyl tail and one polyunsaturated acyl tail (e.g., PG 16:0_18:2, PE 16:0_18:3, and PC 18:0_18:2) (**Figure S8D**). Similarly, TGs enriched in saturated acyl tails (e.g., TG 50:0, TG 52:0, and TG 54:0) were observed in PC1. These data demonstrate an underrepresentation of MUFA acyl tails in the lipidome and support the idea that YTX-7739 treatment of cells significantly decreases the incorporation of MUFAs into complex lipids irrespective of AMPK signaling.

To better understand the changes to the lipidome induced by SCD inhibition in the presence of constitutive AMPK signaling, we examined the lipids positively and negatively contributing to PC2 (**Figures S8E-F**). Positive contributors to PC2 were largely phospholipids containing two saturated acyl tails (e.g., PE 18:0_16:0, PE 16:0_16:0, and PC 18:0_18:0). In contrast, we observed that negative contributors to PC2 (i.e., reflective of AMPK signaling and SCD inhibition) were markedly enriched for phospholipids containing MUFAs (e.g., PG 16:1_18:1, PC 12:0_18:1, and PE 18:0_18:1) or PUFAs (e.g., PE 18:0_20:4, PC 18:2_18:2). These data suggest that AMPK signaling lowers the saturation state of phospholipids in the membrane of SCD-inhibited cells, thereby decreasing engagement of ER stress pathways, lipotoxicity, and cell death.

We also observed several TG and CE species containing SFAs negatively contributing to PC2 (e.g., TG 42:0, CE 18:0, and CE 16:0). These data suggested that AMPK signaling was shuttling SFAs into neutral lipid storage. Bodipy staining of ctrl and AMPK cells confirmed that AMPK signaling increased lipid droplets in GSCs (**Figure 6F**). Cluster analysis of neutral lipids species (e.g., CEs and DGs) showed that AMPK signaling increased the amounts of individual CEs and DGs containing MUFA or PUFA acyl tails (**Figure 6G**). Importantly, SCD inhibition drove the accumulation of SFA containing neutral lipid species only in AMPK GSCs. Likewise, analysis of the desaturation state of TGs revealed a similar pattern where SCD inhibition resulted in the accumulation of saturated or monounsaturated TGs (**Figure 6H**). These data suggested that GSC-AMPK cells treated with YTX-7739 shuttle lipotoxic SFAs (mainly 16:0 and 18:0) into LD to preserve cellular viability. To mimic the intracellular increase of DG levels, we used 1-oleoyl-2-acetyl-glycerol (OAG), a membrane-permeable analog of DG. Supplementing GSCs lines with OAG prevented UPR and lipotoxicity induced by excess palmitate, YTX-7739, or SCD silencing (**Figure 6I-L, S9A-D**). Diacylglycerol O-acyltransferase (DGAT) catalyzes the terminal step converting DG and fatty acyl-CoA into TG, which subsequently form lipid droplets in the cytosol ^42,43^. Pharmacological inhibition of DGAT1 and/or DGAT2, which prevents TG and LD formation in GBM ^44^, reversed OAG-mediated protection against YTX-7739 and C16:0 (**Figure 6K and S9E-F**). Taken together, these results reveal that AMPK activation can overcome DNLS inhibition and resultant lipotoxicity by altering membrane lipid composition and channeling saturated fatty acids into neutral lipids (e.g., CEs and TGs).

## Discussion

This current study explored the therapeutic efficacy of a clinical-stage inhibitor of SCD and the genetic drivers conferring vulnerability to lipogenesis inhibitors in GBM. From a clinical perspective, several of our findings are noteworthy. First, we show that YTX-7739, the first clinical-stage inhibitor of SCD developed to treat synucleinopathies, triggers lipotoxicity in GSCs. We provide evidence supporting the clinical evaluation of this SCD inhibitor in patients diagnosed with GBM. Second, we demonstrate that the activation of MEK/ERK signaling renders GSCs particularly vulnerable to pharmacological inhibitors of lipogenesis. Further, we reveal that the differential GSCs dependence on DNLS is dictated by AMPK activity, thus illustrating a protective role of AMPK and its contribution to metabolic plasticity in GBM.

Preclinical studies in rats and cynomolgus monkeys revealed favorable pharmacokinetic properties of YTX-7739, including bioavailability, low clearance, and a favorable brain-to-plasma ratio of greater than 1 ^13^. Our PD studies in mice bearing GBM tumors showed that YTX-7739 could achieve a comparable reduction in FA desaturation both in the normal brain and in brain tumors, suggesting its ability to penetrate the brain and the brain-tumor barrier effectively. In our preclinical orthotopic GSCs mouse models, treatment with YTX-7739 improved overall survival, particularly in the MGG8-GSCs model. On the other hand, the efficacy of YTX-7739 as monotherapy in the 83 model was less robust. Mice implanted with 83 GSCs presented a short survival of 17-19 days on average, which narrows the therapeutic window (in rats, the maximal reduction of C16 DI was obtained after dosing with YTX-7739 for 7 days ^13^) and limits the overall treatment regimen. For these fast-growing tumors, a higher dose of YTX-7739 or twice-daily (BID) dosing might deplete endogenous pools of SCD products more rapidly, thereby increasing therapeutic efficacy.

Overall, the efficacy of this inhibitor in most animals included in our current study was always transient and, in most cases, failed to eradicate GBM tumors. Therefore, from a clinical perspective, combination therapies with TMZ, radiation, or other agents are more likely to provide a better therapeutic outcome. This is also supported by our observation that inhibiting SCD impairs DNA damage repair through homologous recombination by downregulating RAD51 ^11^. In the highly aggressive 83 model, YTX-7739 with TMZ significantly extended overall survival compared to either monotherapy. In the second MGG8 model (highly sensitive to TMZ), YTX-7739 as monotherapy was equally effective as the standard of care adjuvant chemotherapeutic currently used for GBM treatment.

Identifying genetic drivers of vulnerability to SCD and other DNLS inhibitors in GBM is of major clinical significance. We show that constitutive activation of HRAS/MEK/ERK promotes susceptibility to DNLS inhibitors, while impaired MEK/ERK activity renders GSCs insensitive to the SCD inhibitor. We identified AMPK as a downstream target for MEK/ERK and the primary regulator of sensitivity to DNLS inhibitors. Importantly, our findings reveal an adaptive metabolic reprogramming orchestrated by AMPK, allowing GSCs to overcome DNLS inhibition and maintain proliferation and tumor initiation capacity. We show that GSCs that survive long-term pharmacological inhibition of SCD have decreased MEK/ERK signaling and increased AMPK activity. AMPK activation confers protection against metabolic stress ^38,45^. By employing shotgun lipidomics, we were able to show that AMPK activation allows GSCs to overcome lipotoxicity, sustain proliferation, and retain their in vivo tumor initiation capacity when DNLS is impaired by altering membrane lipid composition and sequestering toxic SFA into neutral lipids. In light of the role of AMPK in promoting a switch from anabolic to catabolic processes ^26^, including the inactivation of ACC ^25^, a defined mechanism for this seemingly contradictory upregulation of MUFA and LDs in GSCs with a constitutively active AMPK remains to be elucidated. AMPK increases FA uptake by enhancing the translocation of the FA transporter CD36 to the plasma membrane ^46^. Therefore, AMPK may enhance CD36-mediated uptake of exogenous FA in GSCs. From that perspective, impeding AMPK activity is likely to maximize the therapeutic efficacy of DNLS-targeting agents. However, this remains a challenge in the absence of highly specific, brain-penetrant inhibitors of AMPK.

Our findings uncover that therapeutic response to DNLS-targeted therapy is contingent on a dynamic shift in molecular signaling, particularly MEK/ERK and AMPK, which, as a nutrient and energy sensor, could be regulated in response to the cellular energetics needs or the biochemical microenvironment of the tumor. This implies bidirectional plasticity where sensitive GSCs could become more resistant to the inhibitor (as seen with 83R and M76R). In contrast, others (such as MGG8) failed to respond to YTX-7739 mediated lipotoxicity in culture yet displayed an excellent therapeutic response following intracranial implantation in mice. Neither the metabolic demands nor the biochemical environment of GBM is faithfully recapitulated under *in vitro* cell culture conditions, and various intrinsic or exogenous stimuli pertinent to the GBM microenvironment can impact MAPK/ERK and AMPK signaling. For instance, the rapid growth of GBM tumors creates hypoxic niches enriched in GSCs ^47,48^, while ERK1/2 activation is triggered by hypoxia ^49,50^. Therefore, GSCs residing in these hypoxic niches may be particularly vulnerable to DNLS inhibition. On the other hand, glucose levels that potently impact AMPK activity and the response to SCD inhibitors, as we have shown, are typically high in the GBM culture medium relative to restricted glucose levels in the brain tumor environment ^51^. Therefore, it is likely that AMPK is activated in glucose-deprived tumors. Other physiological and pathological conditions relevant to GBM including aging (the median age of diagnosis with IDH wild-type GBM is 68-70 years ^52^) and inflammation (pro-inflammatory cytokines are abundant in the GBM microenvironment), suppress AMPK activity ^53^. Further, additional factors, such as diet and caloric intake, are likely to impact the response to SCD inhibitors^16^. Therefore, we propose that the phosphorylation levels of MEK/ERK and AMPK (detected by immunohistochemistry of tumor biopsies) could be predictive biomarkers to guide patient selection and stratification. Our findings should also help tailor treatment paradigms to maximize therapeutic efficacy. For instance, some widely used medicines, such as the anti-inflammatory agent salicylate or anti-diabetic compound Metformin, are potent activators of AMPK ^54,55^ and could be detrimental to the efficacy of YTX-7739 or other DNLS-targeting therapies. On the other hand, despite its potential protumorigenic effect, insulin therapy is commonly used to manage metabolic complications arising from corticosteroid treatment in GBM patients ^40^. Although speculative and requires further validation, it is possible that short-term insulin therapy could maximize the therapeutic efficacy of YTX-7739 in GBM patients, as suggested by our in vitro results.

In conclusion, our current studies highlight metabolic adaptability in GSCs and provide an integrative framework for therapeutic targeting of DNLS in GBM. Our findings support the clinical evaluation of YTX-7739 in patients diagnosed with GBM and reveal predictive biomarkers of therapeutic response to DNLS. We believe that these findings could inform patient selection, drug combination, and clinical trial design to maximize therapeutic efficacy for this morbid and incurable brain cancer.

## MATERIALS AND METHODS

### Primary cells and cell lines

Primary GBM cells were derived from surgical specimens obtained from GBM patients at the Massachusetts General Hospital (provided by Dr. Hiroaki Wakimoto) under the appropriate Institutional Review Board approval (MGG6, MGG8, 70RR) or provided by Dr. Ichiro Nakano (157, 83, 326), Dr. Brent Reynolds (L0, L1, L2) and Dr. Jann Sarkaria at The Brain Tumor PDX National Resource at Mayo Clinic (M76, M12, M120). All GBM cells used in this study have been previously characterized ^56-60^. Cells were expanded as neurospheres and maintained in DMEM/F12 medium supplemented with B27 without vitamin A (1:50; Life Technologies), heparin (2 μg/mL; Sigma Aldrich), human recombinant EGF (20 ng/mL; ABM), and human recombinant bFGF-2 (10ng/mL; ABM). DMEM-F12 w/o L-Glutamine w/o Hepes w/o Glucose (Biowest) was used to control glucose concentrations in the culture medium. NHAs were obtained from Sciencell Research Laboratories and grown in Astrocyte Medium (Sciencell). Human astrocytes transformed with E6/E7/hTERT were provided by Dr. Anna Krichevsky and Dr. Erik Uhlmann and generated as previously described ^24^. Ionizing radiation of GSCs was performed using a ^137^Cs irradiator.

### Lentiviral production

For lentivirus packaging, 293T cells (5 × 10^6^) were seeded in 150 mm plates with Opti-MEM (Gibco; 51985091). After 24h, 30 µL of chloroquine (25 mM; Sigma-Alrich; C6628) was added to the fresh culture medium, and cells were co-transfected with 15 µg plasmids encoding the gene or shRNA of interest, 3.75 µg PMD2.G (a kind gift from Didier Trono, Addgene plasmid 12259) and 11.25 µg psPAX2 (a kind gift from Didier Trono, Addgene plasmid 12260) using PEI (Polysciences, Inc; 23966100) in a 1:3 ratio (total DNA:PEI). At 72 h post-transfection, the conditioned medium was centrifuged at 500 × g for 10 min to remove cell debris. The supernatant was filtered through a 0.45 µm pore size polyethersulfone (PES) filter (Cell Treat; 229749). The filtrate was then centrifuged at 70,000 x g for 90 min at 4°C. Virus pellets were resuspended in 200-500µL PBS/1% BSA, aliquoted, and stored at −80°C.To transduce GSCs or astrocytes, cells (2-5 × 10^5^) were seeded in a six-well plate in the presence of polybrene (10 µg/ml; Sigma-Aldrich; TR-1003G) and 25 µl of lentivirus was added to the well. To generate stable cell lines, cells were subsequently selected using Puromycin (1 µg/ml; Invivogen; ant-pr), G418 (400 µg/ml; Invivogen; ant-gn) or Blasticidin (5-10 µg/ml; Invivogen; ant-bl).

### DNA constructs and Lentiviral vectors

For experiments involving ectopic expression of different genes, a control lentivirus vector pHAGE-CMV-MCS-IRES-ZsGreen (obtained from the DNA Resource Core at Harvard Medical School, HMS) was used as a control vector (Ctrl). pLenti6-AMPK alpha1 (1-312), referred to as AMPKa1, was a gift from Boyi Gan (Addgene plasmid 162131). pLenti-XI-Neo-GST-Constitutively Active AMPK, referred to as AMPKa2, was a gift from Jacob Corn (Addgene plasmid 139843). CSCW-Fluc-IRES-mCherry lentivirus vector carrying an expression cassette for firefly luciferase (Fluc) and mCherry fluorescent protein was used for in vivo imaging. The following shRNA bacterial glycerol stocks: shSCD1 (TRCN0000327814), shAMPKa1 (TRCN0000000861), shAMPKa2 (TRCN0000000861), and non-targeting control shRNA (pLKO.1-puro non-Target shRNA Control; referred to as shCtrl) were obtained from Sigma (MISSION® shRNA Library), amplified, and packaged into lentivirus vectors. GSCs were stably transduced with shRNA lentivirus and, when applicable, selected using puromycin (1 μg/mL), Blasticidin (5-10 μg/mL), or G418 (200-400 μg/mL). Knockdown efficiency was determined using immunoblotting and/or qRT-PCR.

### Preparation of YTX-7739

For cell culture studies, YTX-7739 was solubilized in DMSO. For in vivo studies, YTX-7739 was solubilized by vortexing (1 min), sonication (3-5 min), and heating (3-5 min at 60°C) in 0.5% methylcellulose + 0.2% Tween80 in sterile saline solution.

### Chemical Reagents

The following compounds were obtained from Cayman Chemicals: CAY10566; 4μ8C; NSC95682; SP 600125; DB07268; ND-646; GSK2194069; CP-640186; AZD8330; AZD0364; Azoramide. Salubrinal, Phenylbutyrate, and 1-Oleoyl-2-acetyl-sn-glycerol were obtained from Sigma, Temozolomide, and PF-06424439 from MedChemExpress. A922500 was purchased from Apexbio Technology. Oleate and Palmitate (Cayman Chemicals) were dissolved in DMSO to yield a stock concentration of 0.5-1M. Fatty acids stocks were subsequently complexed to fatty acid-free Bovine Serum Albumin (BSA; 2.5mM; Gold Biotechnology) and added to the culture medium.

### Immunoblot analysis

Cell lysis was performed using RIPA buffer (Boston Bio Products) supplemented with protease and phosphatase inhibitors. Proteins were quantified using the Bradford protein determination assay (Bio-Rad), and 20–40 µg of protein were loaded and resolved on 10% NuPAGE Bis-Tris gels (Life Technologies), transferred to nitrocellulose membranes (Bio-Rad), then incubated with the indicated antibodies. Proteins were detected with SuperSignal West Pico Chemiluminescent Substrate (Pierce). The following antibodies were purchased from Cell Signaling Technologies: SCD (2438); GRP78 (3177); phospho SAPK/JNK (9255); SAPK/JNK (9252); phospho c-Jun (3270); phospho-eIF2α (3597); γ-H2AX (9718); CHOP (2895); PARP (9532); phospho AMPKα (2531); AMPKα (2532); phospho P44/42 MAPK (9101); P44/42 MAPK (4695); Phospho-Acetyl-CoA Carboxylase (11818); Phospho-EGF Receptor (3777); Phospho-AKT (4060); AKT (9272); β-actin (3700); Anti-rabbit IgG, HRP-linked Antibody (7074) and Anti-mouse IgG, HRP-linked Antibody. Phospho IRE1α (AB5700519) and phospho RSK1 (ABS1849) were obtained from Signa-Aldrich. Rad51 (BSM-51402M) and IRE1 (BS-8680R) were obtained from BIOSS Antibodies, GAPDH (sc-47724) from Santa Cruz and eIF2a (ab169528) from AbCam.

### Cell-based Assays

CellTiter-Glo (Promega) was used to measure cell viability. Caspase 3/7 activity was detected using Caspase-Glo 3/7 (Promega). Annexin V/Propidium Iodide staining was performed using Alexa Fluor 488 Annexin V/Dead Cell Apoptosis Kit (Invitrogen). These reagents were used as recommended by the manufacturer. Each data point in the treated samples was normalized to its respective vehicle or pretreatment control for data analysis.

### RNA extraction and PCR

RNA was extracted using the Zymo Quick-RNA MiniPrep kit. RT-PCR was performed with abm 5X AllInOne RT MasterMix. qPCR was run with LUNA universal qPCR master mix with three to four replicates using QuantStudio™ 3 System. All reagents were used as recommended by the manufacturers. Relative mRNA expression was calculated using the 2^-ΔΔCT^ method. qPCR runs included human β-actin and/or HPRT as internal normalization controls. Primer sequences used in this study are listed below. Oligonucleotides were synthesized by the CCIB DNA Core Facility at the Massachusetts General Hospital.

Primer sequences used for qPCR analysis

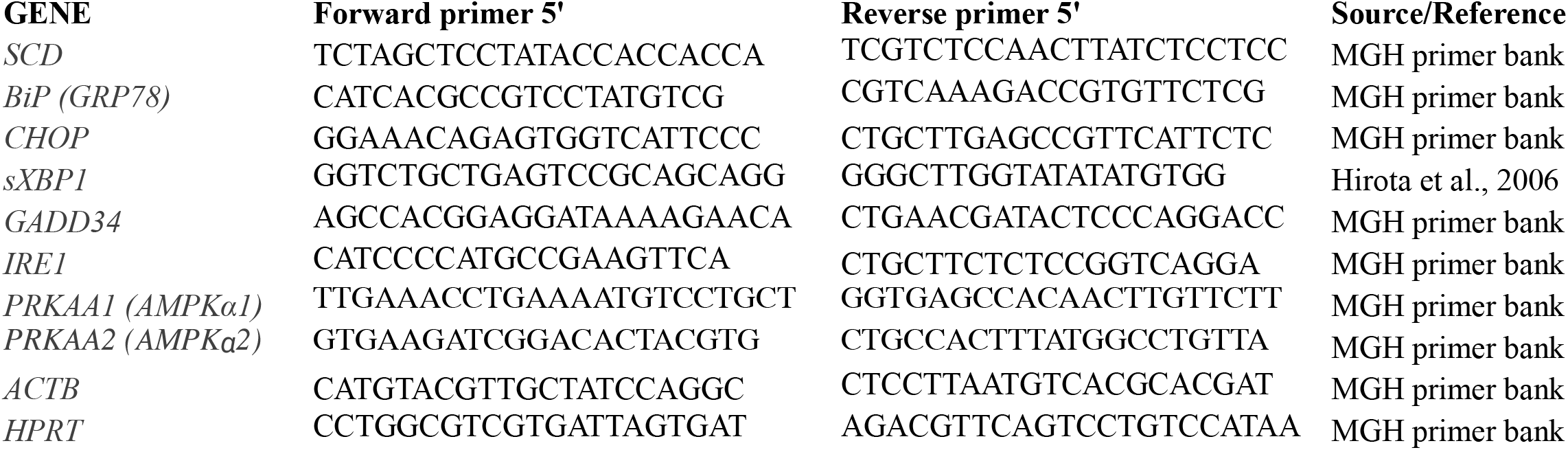

### Mouse Orthotopic Brain Tumor Models

All animal experiments were approved by the Massachusetts General Hospital Subcommittee on Research Animal Care and conformed to the guidelines of the NIH Guide for the Care and Use of Laboratory Animals. GSCs (10-50×10^5^) expressing Firefly luciferase (Fluc) were implanted into the left forebrain of male and female athymic nude mice (1.0 mm anterior and 2.0 mm laterally to bregma at a depth of 2.5 mm from the skull surface) using a stereotactic frame. Brain tumors were detected and monitored over time using Fluc bioluminescence imaging using a Xenogen IVIS 200 imaging system (PerkinElmer) following intraperitoneal (i.p.) injections of D-luciferin (150 mg/kg body weight) (Gold Biotechnology). Signal intensity was quantified with Living Image software 4.3.1 (PerkinElmer). For all in vivo studies, YTX-7739 was administered by i.p. injections. Control groups were administered with 100 µl of vehicle (0.5% methylcellulose + 0.2% Tween80 in sterile saline solution) by i.p. injections. TMZ was dissolved in DMSO (100 mM stock solution) and freshly resuspended in 5% dextrose before i.p. administration in mice. For fatty acid profiling, mice bearing GSCs-brain tumors were treated for 11 days with solvent control or YTX-7739 (30mg/kg). On day 14 after implantation, all mice were sacrificed. Plasma and brain tissue samples (tumor and the contralateral hemisphere) were collected for further analysis.

### Fatty acid profiling

Quantitative measurements on FA for cultured GSCs, plasma, and tissue were performed by OmegaQuant (Sioux Falls, SD) using gas chromatography (GC) with flame ionization detection. Homogenized tissues were extracted with a modified Folch extraction, and portions of the organic layer were used for analysis. Dried cell pellets or tissue extracts were transferred into glass vials and dried in a speed-vac before adding methanol containing 14% boron trifluoride, toluene, methanol; 35:30:35 v/v/v) (Sigma-Aldrich, St. Louis, MO). Vials were vortexed and heated at 100ºC for 45 min. Hexane (EMD Chemicals, USA) and HPLC grade water were added to the cooled samples, followed by vortexing and centrifugation steps to separate layers. An aliquot of the hexane layer was then transferred to a GC vial. GC was carried out using a GC-2010 Gas Chromatograph (Shimadzu Corporation, Columbia, MD) equipped with an SP-2560, 100-m fused silica capillary column (0.25-mm internal diameter, 0.2-um film thickness; Supelco, Bellefonte, PA). A standard mixture of fatty acids characteristic of RBC (GLCOQ-A, NuCheck Prep, Elysian, MN) was used to identify fatty acids and determine individual fatty acid calibration curves. The following 24 fatty acids classes were detected: saturated (14:0, 16:0, 18:0, 20:0, 22:0 24:0); cis monounsaturated (16:1, 18:1, 20:1, 24:1); trans [16:1,18:1*, 18:2*); cis n-6 polyunsaturated (18:2, 18:3, 20:2, 20:3, 20:4, 22:4, 22:5); cis n-3 polyunsaturated (18:3, 20:5, 22:5, 22:6). The fatty acid composition was expressed as a percent of the total identified fatty acids. SCD-mediated desaturation was determined by measuring the Desaturation index (DI) as follows: C16 DI= C16:1n-7/C16:0; C18 DI = C18:1n-9/C18:0.

### Shotgun Lipidomics Analysis

83 cells were seeded at a density of 1×10^5^ cells/well in a 6-well plate and treated with DMSO (Control) or YTX-7739 (1 µM) in four replicates. At 72h after treatment, cells were counted and collected for lipid extraction. Shotgun lipidomics analysis was performed as previously described ^61^. Briefly, after the addition of an internal standard mixture consisting of 70 lipid standards across 17 subclasses (AB Sciex, 5040156, Avanti 330827, Avanti 330830, Avanti 330828, Avanti 791642), and two successive extractions, organic layers were vacuum dried in a Thermo SpeedVac SPD300DDA machine, using ramp setting 4 at 35ºC for 45 minutes with a total run time of 90 minutes. Samples were then resuspended in methanol/dichloromethane (1:1 ratio) with 10 mM Ammonium Acetate and transferred to robovials (Thermo 10800107) for subsequent lipidomics analysis. Samples were analyzed by direct infusion using the Sciex 5500 with Differential Mobility Device (DMS) (comparable to Sciex Lipidyzer platform) for quantitative measurement of 1450 lipid species belonging to 17 subclasses. The DMS was tuned with EquiSPLASH LIPIDOMIX (Avanti 330731). Instrument settings, MRM lists, and data analysis methods are available ^62^. Quantitative values of lipid species were normalized to cell counts. Heatmaps and PCA plots were generated using Clustvis ^63^. The color scale on heatmaps refers to the row Z score. Nipals PCA was utilized to calculate principal components (PC) on PCA. X and Y axis show PC1 and PC2, which refer to the percentage of the total variance, respectively.

### Immunocytochemistry and Lipid Droplet staining

Cells were fixed with cold acetone for 20 min. Cells were mounted on slides, dried, permeabilized with 0.1% Triton X-100, and blocked with 5% BSA for 1h at room temperature. Cells were then incubated overnight at 4°C with p-ERK1/2 antibody (1:400). Fluorophore-conjugated secondary antibodies (Life Technologies, 1:100) were added for 1h. Nuclei were counterstained with DAPI (Life Technologies), mounted on a microscope slide, and analyzed by fluorescence microscopy. BODIPY 493/503 dissolved in ethanol (1mg/ml stock) was used to stain lipid droplets. Cells were washed 2 times with PBS and incubated with BODIPY 493/503 at 1µg/ml for 30 minutes before mounting and analysis with fluorescent microscopy. The number of lipid droplets per cell was manually counted in a total of 100 cells for each condition.

### Bioluminescence BLRR assay to monitor DNA damage repair

The BLRR system ^23^ was used to track DNA damage repair through Homology directed repair (HDR) and non-homologous end-joining (NHEJ) as previously described ^64^. In brief, GSCs were sequentially transduced to express the three components of this reporter: BLRR, trGluc, and I-SceI. To detect BLRR activity, GSCs were seeded in a 96-well plate and treated with YTX-7739. On day 3 post-treatment, an aliquot of the conditioned medium (25µl) was collected and transferred into a 96-well white plate to measure bioluminescence using a multimode reader (Biotek). Gaussia luciferase (Gluc; HDR) and Vargula luciferase (Vluc; NHEJ) activities were measured by adding coelenterazine (20µM; Nanolight) and Vargulin (5ng/mL; Nanolight), respectively. All bioluminescence results were normalized to cell viability measured using CellTiter-Glo.

### Statistical Analysis

All cell culture experiments consisted of at least three independent replicates and were repeated at least 3 times. The results are presented as individual values or mean ± standard deviation (SD) unless stated otherwise. Statistical significance was determined using GraphPad Prism 9 (GraphPad Software). For two groups comparison, statistical significance was calculated using the two-tailed Student’s t-test when data were normally distributed and the Mann-Whitney test when data were not normally distributed. For multiple groups comparison, p values were generated using one-way ANOVA followed by Dunnett’s multiple comparisons test when data were normally distributed and the Kruskal-Wallis test followed by Dunn’s multiple comparison test when data were not normally distributed to compare different groups to controls. Normality was tested using the Shapiro-Wilk test. Animal survival curves were analyzed for statistical significance using a log-rank test (Mantel-Cox), and survival curves were plotted as Kaplan-Meier using GraphPad Prism 9. P values of 0.05 or less were considered significant. The group size was determined based on preliminary experiments, and no statistical method was used to determine the sample size. Graphs were generated using GraphPad Prism 9 or R package version 0.4.0. https://CRAN.R-project.org/package=ggpubr. Data analysis for lipidomics studies was conducted using R. Schematic figures were created with BioRender.com.

## ACKNOWLEDGMENTS

We are grateful to Drs. Hiroaki Wakimoto (MGH), the Brain Tumor PDX National Resource at Mayo Clinic, Ichiro Nakano (University of Alabama Birmingham), and Brent Reynolds (University of Florida) for providing primary GBM cells; Dr. Anna Krichevsky and Dr. Erik Uhlmann for providing HA-NT cells. We thank Dr. Fei Tian for his technical assistance.

## REFERENCES

1. Badr, C.E., Silver, D.J., Siebzehnrubl, F.A. & Deleyrolle, L.P. Metabolic heterogeneity and adaptability in brain tumors. Cellular and molecular life sciences : CMLS 77, 5101–5119 (2020).

2. Koundouros, N. & Poulogiannis, G. Reprogramming of fatty acid metabolism in cancer. British journal of cancer 122, 4–22 (2020).

3. Hoy, A.J., Nagarajan, S.R. & Butler, L.M. Tumour fatty acid metabolism in the context of therapy resistance and obesity. Nature reviews. Cancer 21, 753–766 (2021).

4. Beloribi-Djefaflia, S., Vasseur, S. & Guillaumond, F. Lipid metabolic reprogramming in cancer cells. Oncogenesis 5, e189 (2016).

5. Verhaak, R.G., et al. Integrated genomic analysis identifies clinically relevant subtypes of glioblastoma characterized by abnormalities in PDGFRA, IDH1, EGFR, and NF1. Cancer cell 17, 98–110 (2010).

6. Brennan, C.W., et al. The somatic genomic landscape of glioblastoma. Cell 155, 462–477 (2013).

7. Jeuken, J., van den Broecke, C., Gijsen, S., Boots-Sprenger, S. & Wesseling, P. RAS/RAF pathway activation in gliomas: the result of copy number gains rather than activating mutations. Acta Neuropathol 114, 121–133 (2007).

8. Kim, J.Y., Kim, Y.J., Lee, S. & Park, J.H. The critical role of ERK in death resistance and invasiveness of hypoxia-selected glioblastoma cells. BMC Cancer 9, 27 (2009).

9. Garcia, D. & Shaw, R.J. AMPK: Mechanisms of Cellular Energy Sensing and Restoration of Metabolic Balance. Mol Cell 66, 789–800 (2017).

10. Eberle, D., Hegarty, B., Bossard, P., Ferre, P. & Foufelle, F. SREBP transcription factors: master regulators of lipid homeostasis. Biochimie 86, 839–848 (2004).

11. Pinkham, K., et al. Stearoyl CoA Desaturase Is Essential for Regulation of Endoplasmic Reticulum Homeostasis and Tumor Growth in Glioblastoma Cancer Stem Cells. Stem Cell Reports 12, 712–727 (2019).

12. Oatman, N., et al. Mechanisms of stearoyl CoA desaturase inhibitor sensitivity and acquired resistance in cancer. Sci Adv 7(2021).

13. Tardiff, D.F., et al. Non-clinical Pharmacology of YTX-7739: a Clinical Stage Stearoyl-CoA Desaturase Inhibitor Being Developed for Parkinson’s Disease. Mol Neurobiol (2022).

14. Marciniak, S.J., et al. CHOP induces death by promoting protein synthesis and oxidation in the stressed endoplasmic reticulum. Genes & development 18, 3066–3077 (2004).

15. Zinszner, H., et al. CHOP is implicated in programmed cell death in response to impaired function of the endoplasmic reticulum. Genes & development 12, 982–995 (1998).

16. Szegezdi, E., Logue, S.E., Gorman, A.M. & Samali, A. Mediators of endoplasmic reticulum stress-induced apoptosis. EMBO reports 7, 880–885 (2006).

17. Lieber, M.R. The mechanism of double-strand DNA break repair by the nonhomologous DNA end-joining pathway. Annu Rev Biochem 79, 181–211 (2010).

18. Chang, H.H.Y., Pannunzio, N.R., Adachi, N. & Lieber, M.R. Non-homologous DNA end joining and alternative pathways to double-strand break repair. Nat Rev Mol Cell Biol 18, 495–506 (2017).

19. Maacke, H., et al. Over-expression of wild-type Rad51 correlates with histological grading of invasive ductal breast cancer. Int J Cancer 88, 907–913 (2000).

20. Gil Del Alcazar, C.R., Todorova, P.K., Habib, A.A., Mukherjee, B. & Burma, S. Augmented HR Repair Mediates Acquired Temozolomide Resistance in Glioblastoma. Molecular cancer research : MCR 14, 928–940 (2016).

21. Short, S.C., et al. Rad51 inhibition is an effective means of targeting DNA repair in glioma models and CD133+ tumor-derived cells. Neuro-oncology 13, 487–499 (2011).

22. King, H.O., et al. RAD51 Is a Selective DNA Repair Target to Radiosensitize Glioma Stem Cells. Stem Cell Reports 8, 125–139 (2017).

23. Chien, J.C., et al. A multiplexed bioluminescent reporter for sensitive and non-invasive tracking of DNA double strand break repair dynamics in vitro and in vivo. Nucleic acids research 48, e100 (2020).

24. Sonoda, Y., et al. Formation of intracranial tumors by genetically modified human astrocytes defines four pathways critical in the development of human anaplastic astrocytoma. Cancer research 61, 4956–4960 (2001).

25. Yeh, L.A., Lee, K.H. & Kim, K.H. Regulation of rat liver acetyl-CoA carboxylase. Regulation of phosphorylation and inactivation of acetyl-CoA carboxylase by the adenylate energy charge. The Journal of biological chemistry 255, 2308–2314 (1980).

26. Trefts, E. & Shaw, R.J. AMPK: restoring metabolic homeostasis over space and time. Mol Cell 81, 3677–3690 (2021).

27. Ma, Y., et al. Antithrombin up-regulates AMP-activated protein kinase signalling during myocardial ischaemia/reperfusion injury. Thromb Haemost 113, 338–349 (2015).

28. Filippov, S., et al. ETC-1002 regulates immune response, leukocyte homing, and adipose tissue inflammation via LKB1-dependent activation of macrophage AMPK. Journal of lipid research 54, 2095–2108 (2013).

29. Hardie, D.G. AMPK--sensing energy while talking to other signaling pathways. Cell Metab 20, 939–952 (2014).

30. Sanduja, S., et al. AMPK promotes tolerance to Ras pathway inhibition by activating autophagy. Oncogene 35, 5295–5303 (2016).

31. Martin, M.J., Carling, D. & Marais, R. Taking the stress out of melanoma. Cancer cell 15, 163–164 (2009).

32. Bhagwat, S.V., et al. ERK Inhibitor LY3214996 Targets ERK Pathway-Driven Cancers: A Therapeutic Approach Toward Precision Medicine. Molecular cancer therapeutics 19, 325–336 (2020).

33. Morris, E.J., et al. Discovery of a novel ERK inhibitor with activity in models of acquired resistance to BRAF and MEK inhibitors. Cancer discovery 3, 742–750 (2013).

34. Pratilas, C.A. & Solit, D.B. Targeting the mitogen-activated protein kinase pathway: physiological feedback and drug response. Clinical cancer research : an official journal of the American Association for Cancer Research 16, 3329–3334 (2010).

35. Liu, X., et al. LncRNA NBR2 engages a metabolic checkpoint by regulating AMPK under energy stress. Nature cell biology 18, 431–442 (2016).

36. Liang, J.R., et al. A Genome-wide ER-phagy Screen Highlights Key Roles of Mitochondrial Metabolism and ER-Resident UFMylation. Cell 180, 1160–1177 e1120 (2020).

37. Dasgupta, B. & Seibel, W. Compound C/Dorsomorphin: Its Use and Misuse as an AMPK Inhibitor. Methods Mol Biol 1732, 195–202 (2018).

38. Chhipa, R.R., et al. AMP kinase promotes glioblastoma bioenergetics and tumour growth. Nature cell biology 20, 823–835 (2018).

39. Laderoute, K.R., et al. 5’-AMP-activated protein kinase (AMPK) is induced by low-oxygen and glucose deprivation conditions found in solid-tumor microenvironments. Molecular and cellular biology 26, 5336–5347 (2006).

40. Gong, Y., et al. Insulin-mediated signaling promotes proliferation and survival of glioblastoma through Akt activation. Neuro-oncology 18, 48–57 (2016).

41. Saltiel, A.R. Insulin signaling in health and disease. The Journal of clinical investigation 131(2021).

42. Greenberg, A.S., et al. The role of lipid droplets in metabolic disease in rodents and humans. The Journal of clinical investigation 121, 2102–2110 (2011).

43. Yen, C.L., Stone, S.J., Koliwad, S., Harris, C. & Farese, R.V., Jr. Thematic review series: glycerolipids. DGAT enzymes and triacylglycerol biosynthesis. Journal of lipid research 49, 2283–2301 (2008).

44. Cheng, X., et al. Targeting DGAT1 Ameliorates Glioblastoma by Increasing Fat Catabolism and Oxidative Stress. Cell Metab 32, 229–242 e228 (2020).

45. Chae, Y.C., et al. Control of tumor bioenergetics and survival stress signaling by mitochondrial HSP90s. Cancer cell 22, 331–344 (2012).

46. Habets, D.D., et al. Crucial role for LKB1 to AMPKalpha2 axis in the regulation of CD36-mediated long-chain fatty acid uptake into cardiomyocytes. Biochimica et biophysica acta 1791, 212–219 (2009).

47. Li, Z., et al. Hypoxia-inducible factors regulate tumorigenic capacity of glioma stem cells. Cancer cell 15, 501–513 (2009).

48. Seidel, S., et al. A hypoxic niche regulates glioblastoma stem cells through hypoxia inducible factor 2 alpha. Brain 133, 983–995 (2010).

49. Li, L., et al. The requirement of extracellular signal-related protein kinase pathway in the activation of hypoxia inducible factor 1 alpha in the developing rat brain after hypoxia-ischemia. Acta Neuropathol 115, 297–303 (2008).

50. Minet, E., et al. ERK activation upon hypoxia: involvement in HIF-1 activation. FEBS letters 468, 53–58 (2000).

51. Flavahan, W.A., et al. Brain tumor initiating cells adapt to restricted nutrition through preferential glucose uptake. Nat Neurosci 16, 1373–1382 (2013).

52. Ostrom, Q.T., et al. CBTRUS Statistical Report: Primary Brain and Other Central Nervous System Tumors Diagnosed in the United States in 2013-2017. Neuro-oncology 22, iv1–iv96 (2020).

53. Jeon, S.M. Regulation and function of AMPK in physiology and diseases. Exp Mol Med 48, e245 (2016).

54. Hawley, S.A., et al. The ancient drug salicylate directly activates AMP-activated protein kinase. Science 336, 918–922 (2012).

55. Zhou, G., et al. Role of AMP-activated protein kinase in mechanism of metformin action. The Journal of clinical investigation 108, 1167–1174 (2001).

56. Wakimoto, H., et al. Maintenance of primary tumor phenotype and genotype in glioblastoma stem cells. Neuro-oncology 14, 132–144 (2012).

57. Tanaka, S., et al. Genetically distinct glioma stem-like cell xenografts established from paired glioblastoma samples harvested before and after molecularly targeted therapy. Scientific reports 9, 139 (2019).

58. Mao, P., et al. Mesenchymal glioma stem cells are maintained by activated glycolytic metabolism involving aldehyde dehydrogenase 1A3. Proceedings of the National Academy of Sciences of the United States of America 110, 8644–8649 (2013).

59. Siebzehnrubl, F.A., et al. The ZEB1 pathway links glioblastoma initiation, invasion and chemoresistance. EMBO Mol Med 5, 1196–1212 (2013).

60. Vaubel, R.A., et al. Genomic and Phenotypic Characterization of a Broad Panel of Patient-Derived Xenografts Reflects the Diversity of Glioblastoma. Clinical cancer research : an official journal of the American Association for Cancer Research 26, 1094–1104 (2020).

61. Hsieh, W.Y., Williams, K.J., Su, B. & Bensinger, S.J. Profiling of mouse macrophage lipidome using direct infusion shotgun mass spectrometry. STAR Protoc 2, 100235 (2021).

62. Su, B., et al. A DMS Shotgun Lipidomics Workflow Application to Facilitate High-Throughput, Comprehensive Lipidomics. J Am Soc Mass Spectrom 32, 2655–2663 (2021).

63. Metsalu, T. & Vilo, J. ClustVis: a web tool for visualizing clustering of multivariate data using Principal Component Analysis and heatmap. Nucleic acids research 43, W566–570 (2015).

64. Chien, J.C., Badr, C.E. & Lai, C.P. Multiplexed bioluminescence-mediated tracking of DNA double-strand break repairs in vitro and in vivo. Nature protocols 16, 3933–3953 (2021).

