## Supplemental Figures for "Divergent MEK/ERK and AMPK signaling dictate lipogenic plasticity and dependence on fatty acid synthesis in Glioblastoma"

### Slide 1
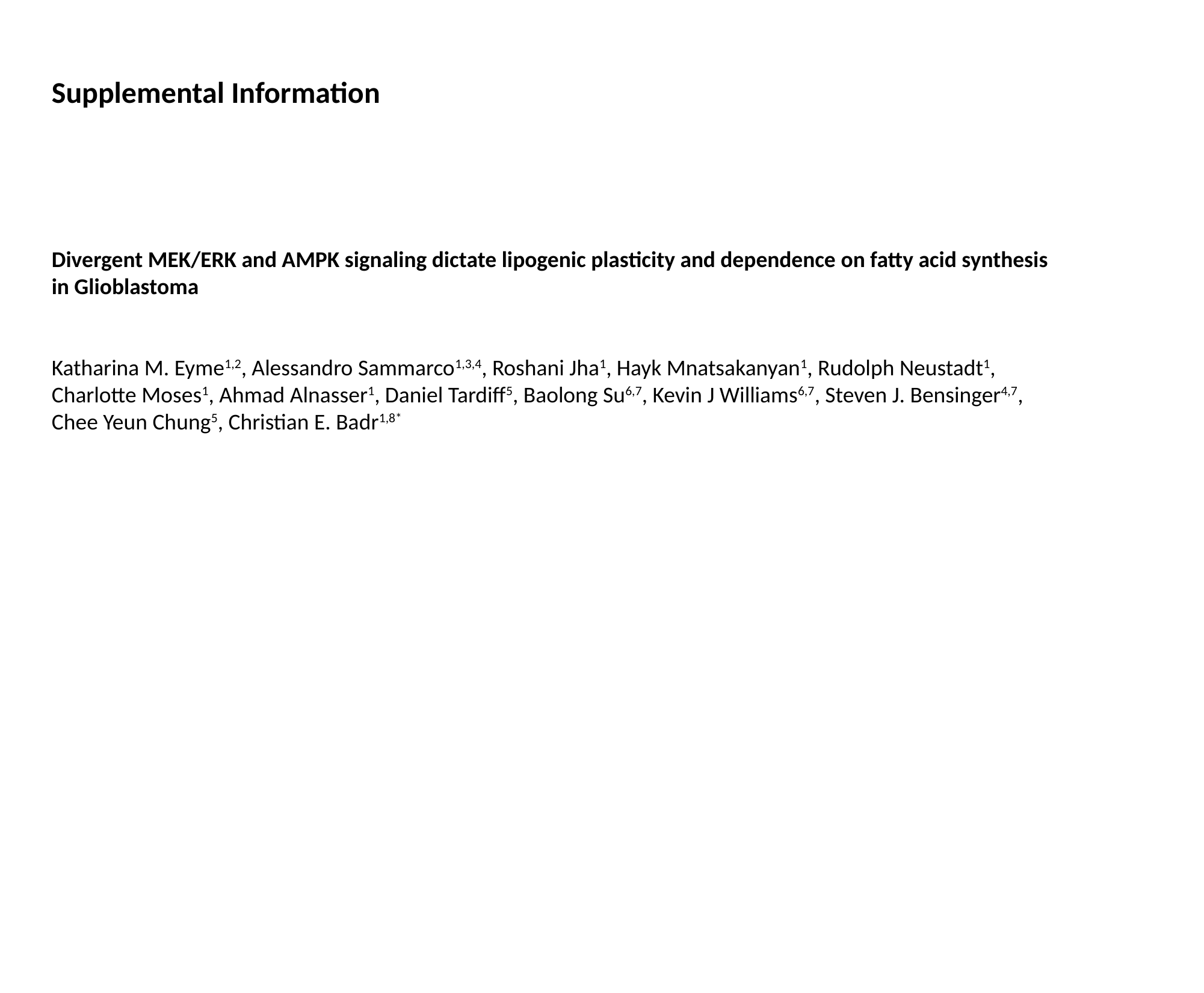

Supplemental Information
Divergent MEK/ERK and AMPK signaling dictate lipogenic plasticity and dependence on fatty acid synthesis in Glioblastoma
Katharina M. Eyme^1,2^, Alessandro Sammarco^1,3,4^, Roshani Jha^1^, Hayk Mnatsakanyan^1^, Rudolph Neustadt^1^, Charlotte Moses^1^, Ahmad Alnasser^1^, Daniel Tardiff^5^, Baolong Su^6,7^, Kevin J Williams^6,7^, Steven J. Bensinger^4,7^, Chee Yeun Chung^5^, Christian E. Badr^1,8*^

### Slide 2
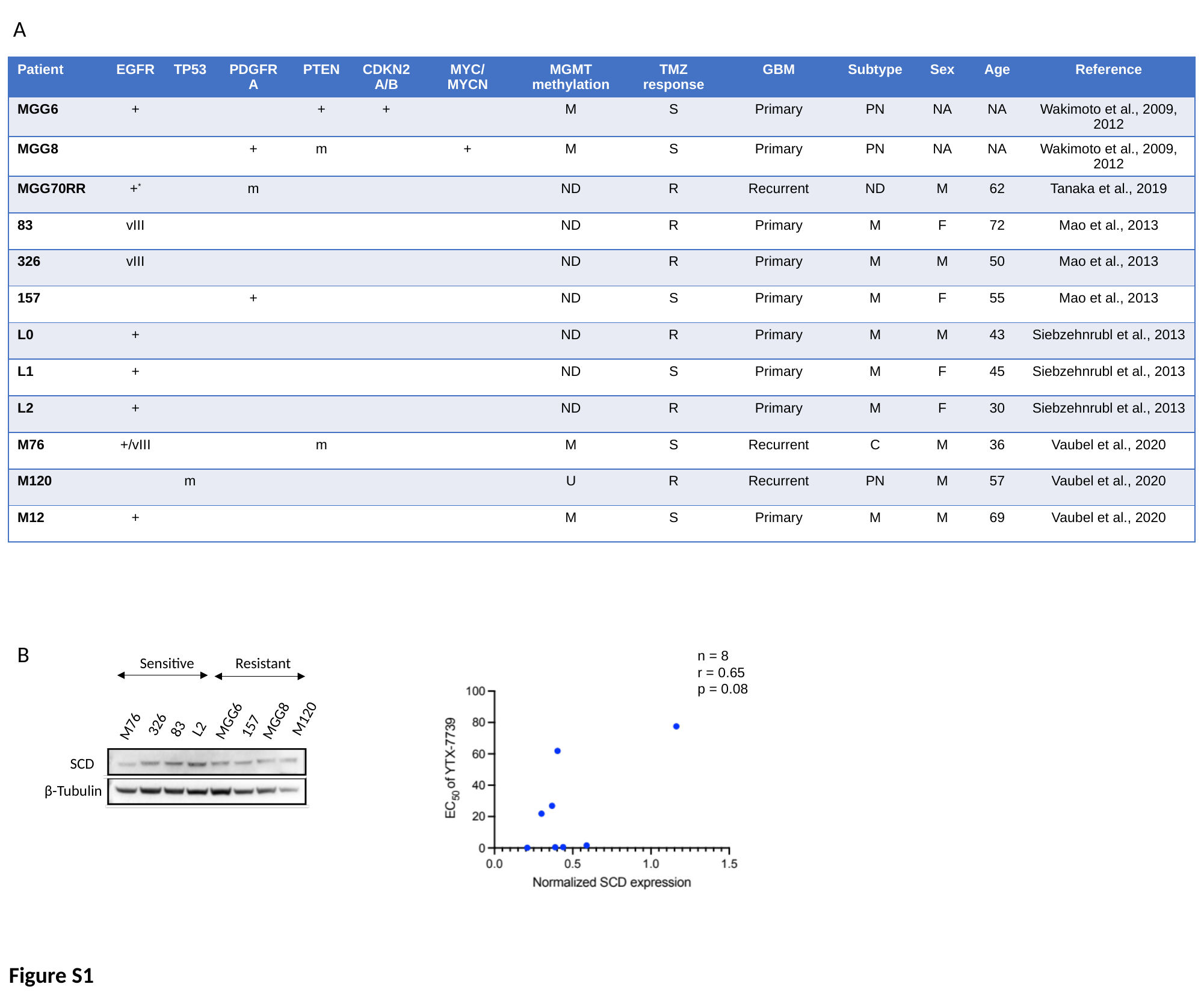

A
| Patient | EGFR | TP53 | PDGFRA | PTEN | CDKN2 A/B | MYC/ MYCN | MGMT methylation | TMZ response | GBM | Subtype | Sex | Age | Reference |
| --- | --- | --- | --- | --- | --- | --- | --- | --- | --- | --- | --- | --- | --- |
| MGG6 | + | | | + | + | | M | S | Primary | PN | NA | NA | Wakimoto et al., 2009, 2012 |
| MGG8 | | | + | m | | + | M | S | Primary | PN | NA | NA | Wakimoto et al., 2009, 2012 |
| MGG70RR | +^\*^ | | m | | | | ND | R | Recurrent | ND | M | 62 | Tanaka et al., 2019 |
| 83 | vIII | | | | | | ND | R | Primary | M | F | 72 | Mao et al., 2013 |
| 326 | vIII | | | | | | ND | R | Primary | M | M | 50 | Mao et al., 2013 |
| 157 | | | + | | | | ND | S | Primary | M | F | 55 | Mao et al., 2013 |
| L0 | + | | | | | | ND | R | Primary | M | M | 43 | Siebzehnrubl et al., 2013 |
| L1 | + | | | | | | ND | S | Primary | M | F | 45 | Siebzehnrubl et al., 2013 |
| L2 | + | | | | | | ND | R | Primary | M | F | 30 | Siebzehnrubl et al., 2013 |
| M76 | +/vIII | | | m | | | M | S | Recurrent | C | M | 36 | Vaubel et al., 2020 |
| M120 | | m | | | | | U | R | Recurrent | PN | M | 57 | Vaubel et al., 2020 |
| M12 | + | | | | | | M | S | Primary | M | M | 69 | Vaubel et al., 2020 |
B
n = 8
r = 0.65
p = 0.08
Sensitive Resistant
326
M76
157
MGG6
MGG8
M120
L2
83
SCD
β-Tubulin
Figure S1

### Slide 3
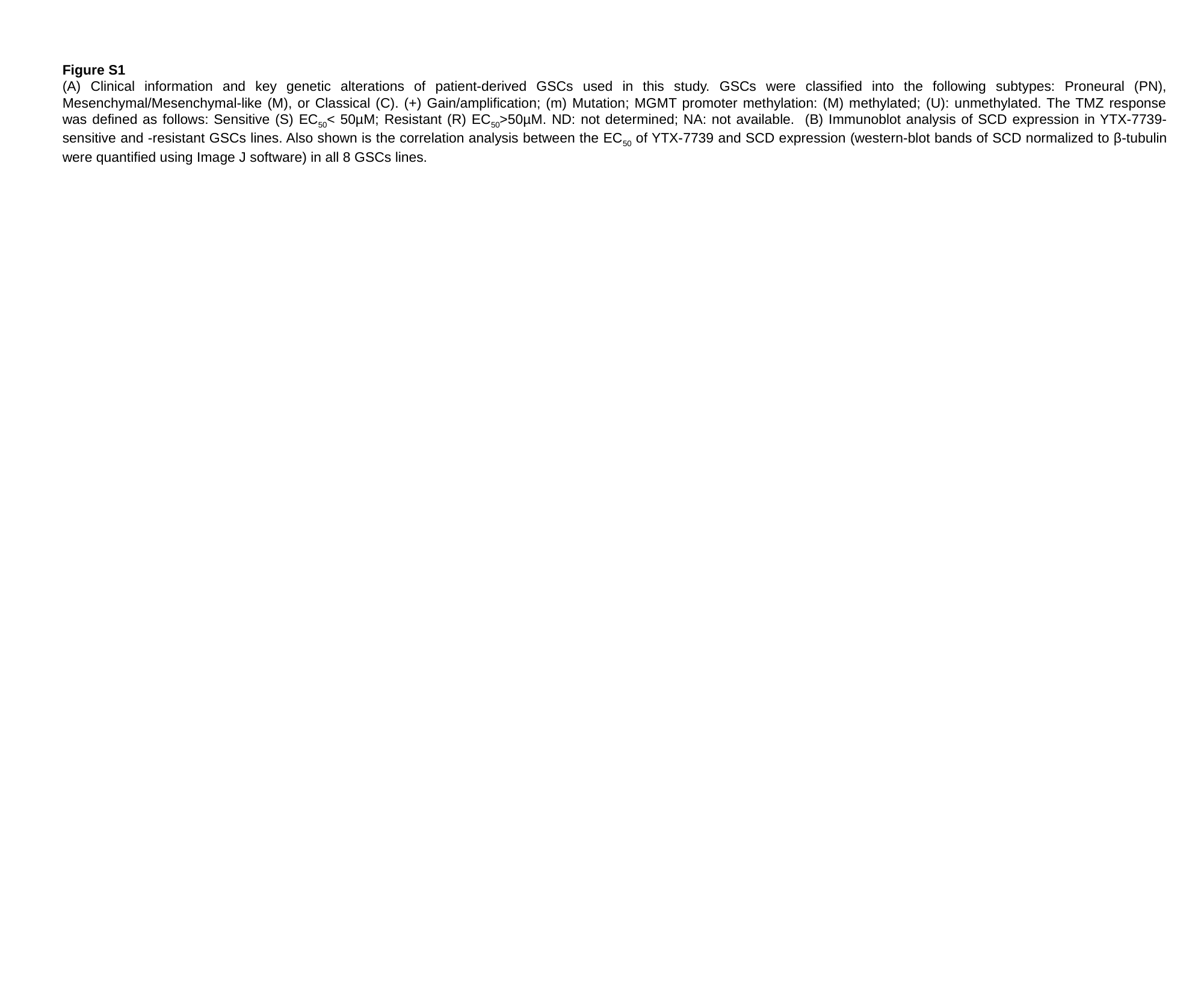

Figure S1
(A) Clinical information and key genetic alterations of patient-derived GSCs used in this study. GSCs were classified into the following subtypes: Proneural (PN), Mesenchymal/Mesenchymal-like (M), or Classical (C). (+) Gain/amplification; (m) Mutation; MGMT promoter methylation: (M) methylated; (U): unmethylated. The TMZ response was defined as follows: Sensitive (S) EC_50_< 50µM; Resistant (R) EC_50_>50µM. ND: not determined; NA: not available. (B) Immunoblot analysis of SCD expression in YTX-7739-sensitive and -resistant GSCs lines. Also shown is the correlation analysis between the EC_50_ of YTX-7739 and SCD expression (western-blot bands of SCD normalized to β-tubulin were quantified using Image J software) in all 8 GSCs lines.

### Slide 4
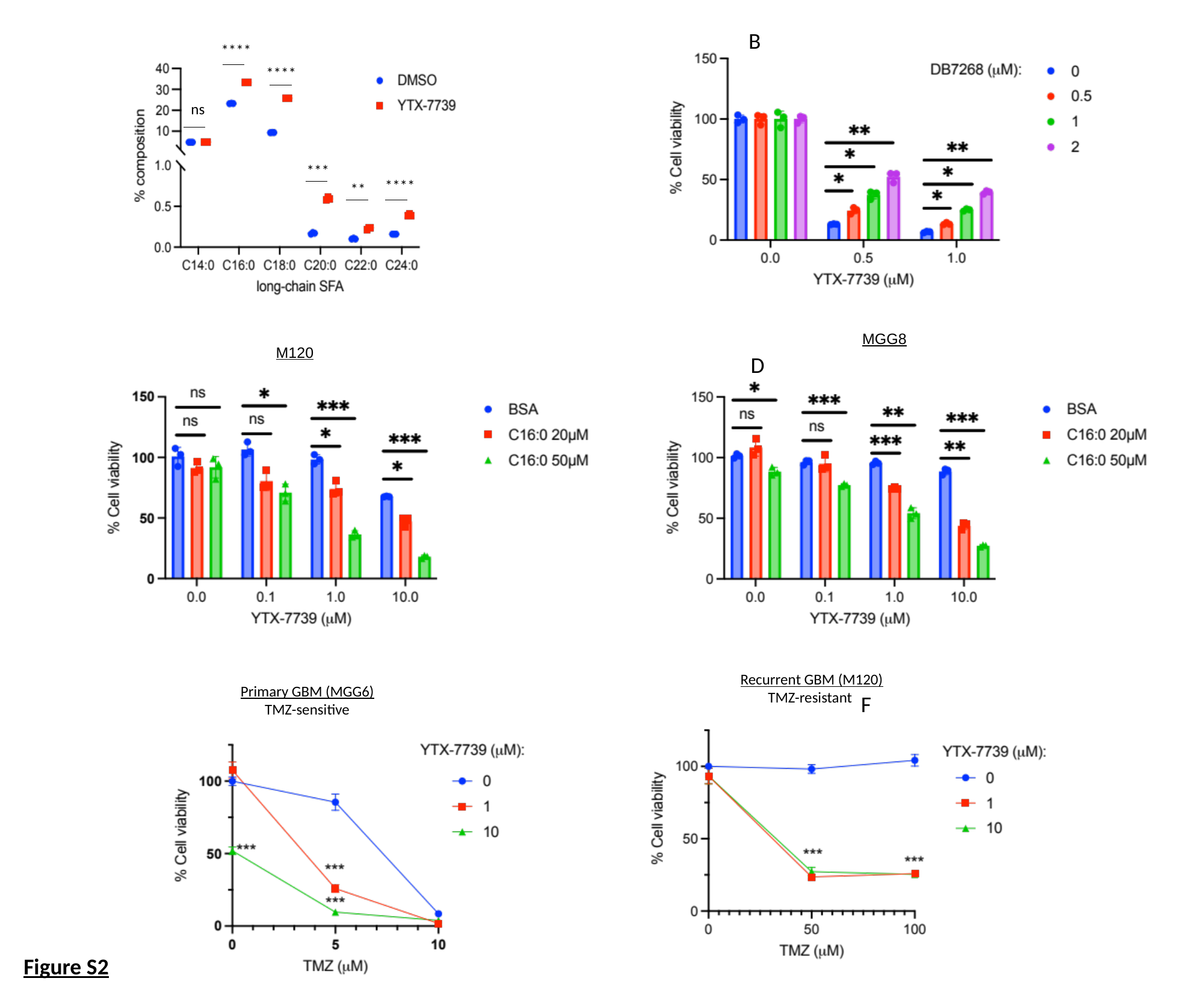

A						 B
****
****
ns
***
****
**
MGG8
M120
C												 D
Recurrent GBM (M120)
 TMZ-resistant
Primary GBM (MGG6)
 TMZ-sensitive
E											 F
Figure S2

### Slide 5
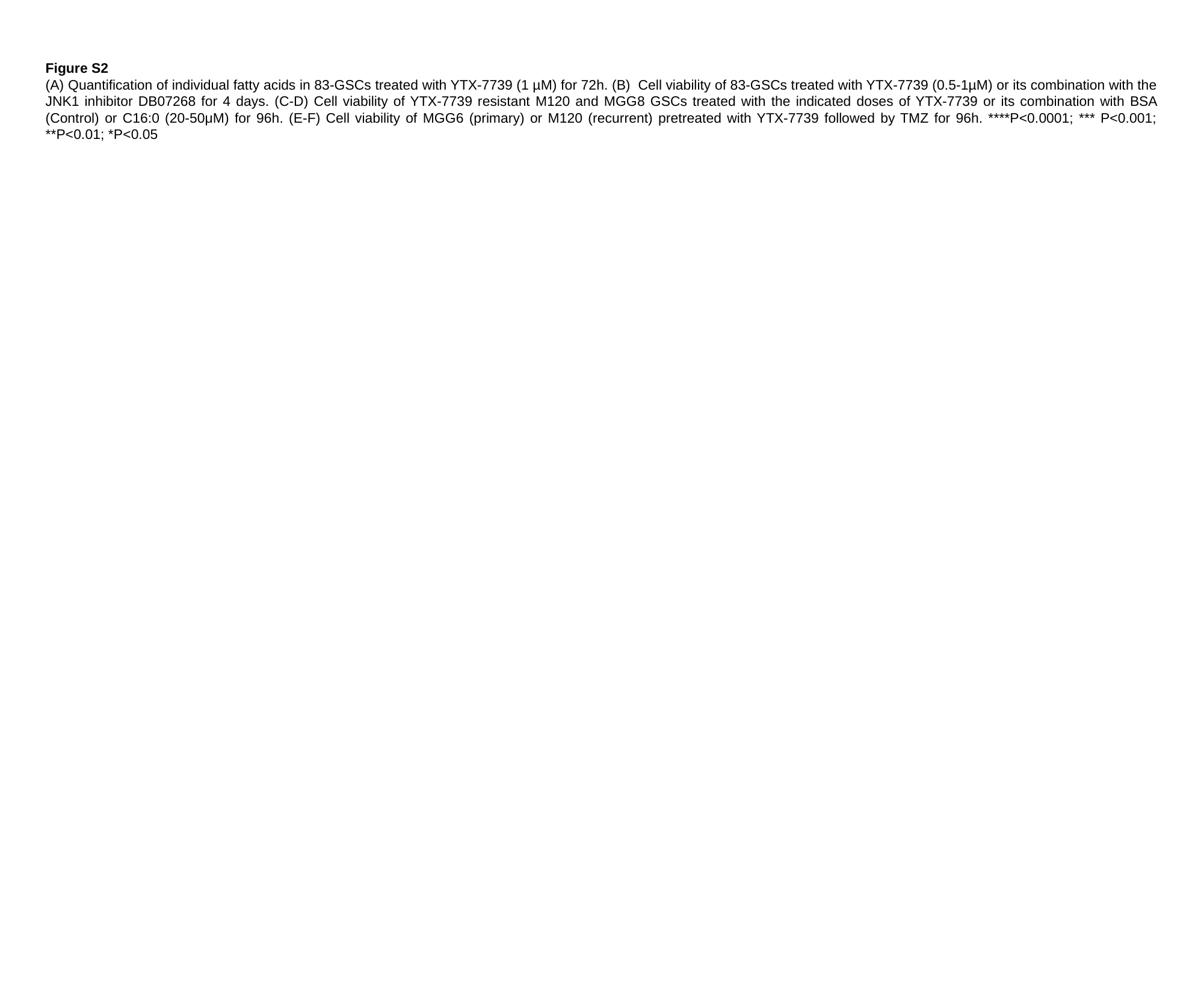

Figure S2
(A) Quantification of individual fatty acids in 83-GSCs treated with YTX-7739 (1 µM) for 72h. (B) Cell viability of 83-GSCs treated with YTX-7739 (0.5-1µM) or its combination with the JNK1 inhibitor DB07268 for 4 days. (C-D) Cell viability of YTX-7739 resistant M120 and MGG8 GSCs treated with the indicated doses of YTX-7739 or its combination with BSA (Control) or C16:0 (20-50μM) for 96h. (E-F) Cell viability of MGG6 (primary) or M120 (recurrent) pretreated with YTX-7739 followed by TMZ for 96h. ****P<0.0001; *** P<0.001; **P<0.01; *P<0.05

### Slide 6
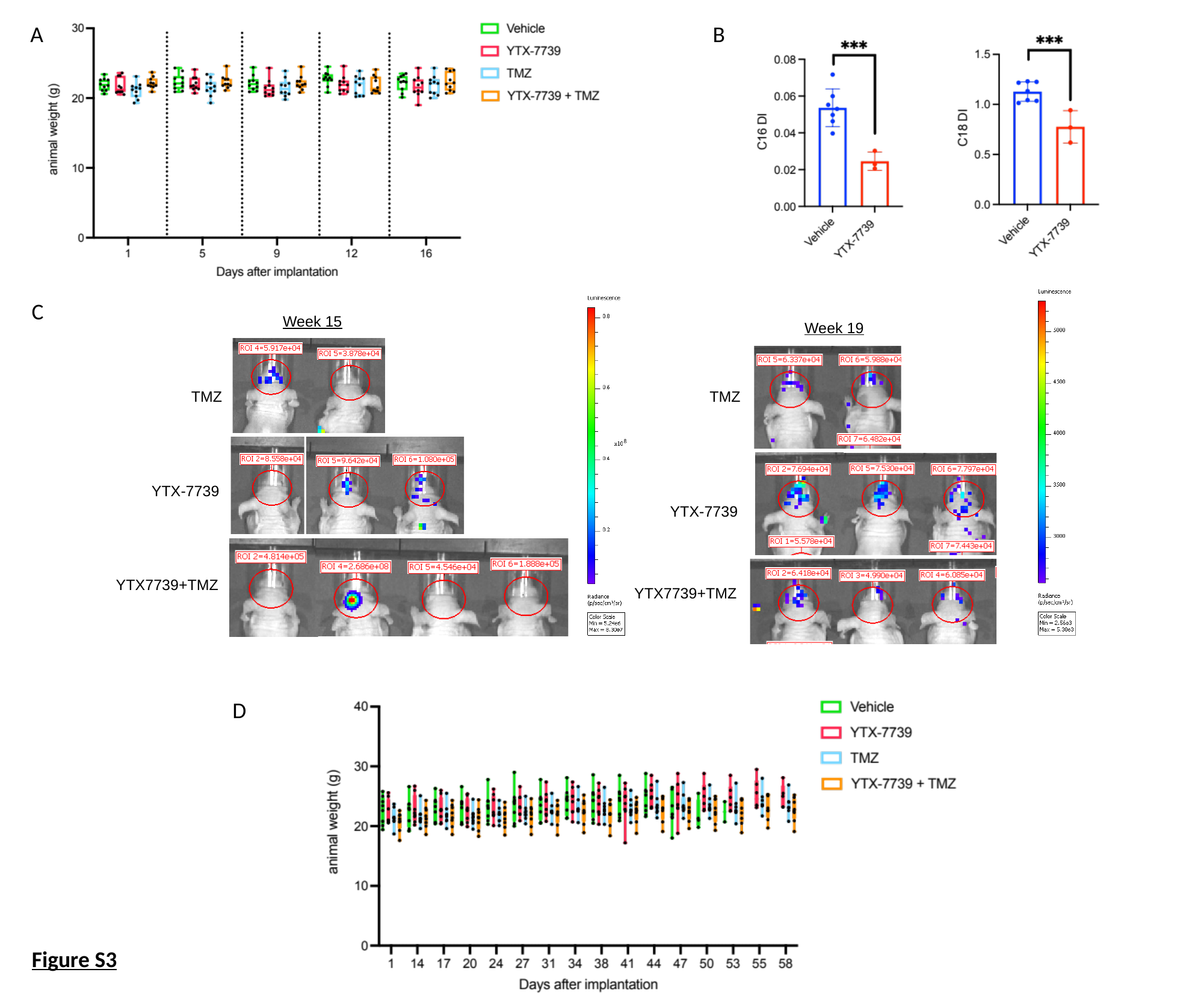

A
B
Week 15
Week 19
TMZ
TMZ
YTX-7739
YTX-7739
YTX7739+TMZ
YTX7739+TMZ
C
D
Figure S3

### Slide 7
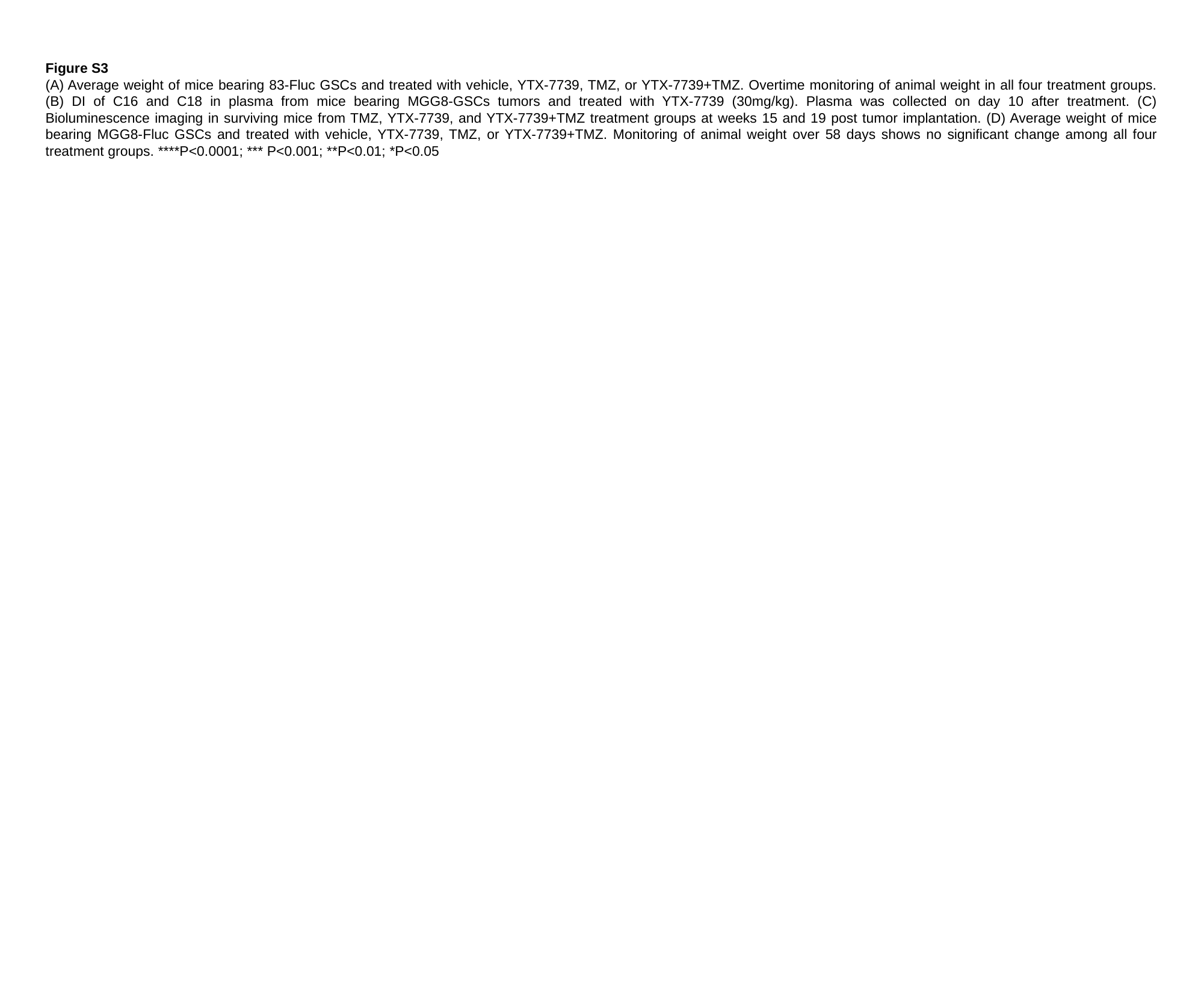

Figure S3
(A) Average weight of mice bearing 83-Fluc GSCs and treated with vehicle, YTX-7739, TMZ, or YTX-7739+TMZ. Overtime monitoring of animal weight in all four treatment groups. (B) DI of C16 and C18 in plasma from mice bearing MGG8-GSCs tumors and treated with YTX-7739 (30mg/kg). Plasma was collected on day 10 after treatment. (C) Bioluminescence imaging in surviving mice from TMZ, YTX-7739, and YTX-7739+TMZ treatment groups at weeks 15 and 19 post tumor implantation. (D) Average weight of mice bearing MGG8-Fluc GSCs and treated with vehicle, YTX-7739, TMZ, or YTX-7739+TMZ. Monitoring of animal weight over 58 days shows no significant change among all four treatment groups. ****P<0.0001; *** P<0.001; **P<0.01; *P<0.05

### Slide 8
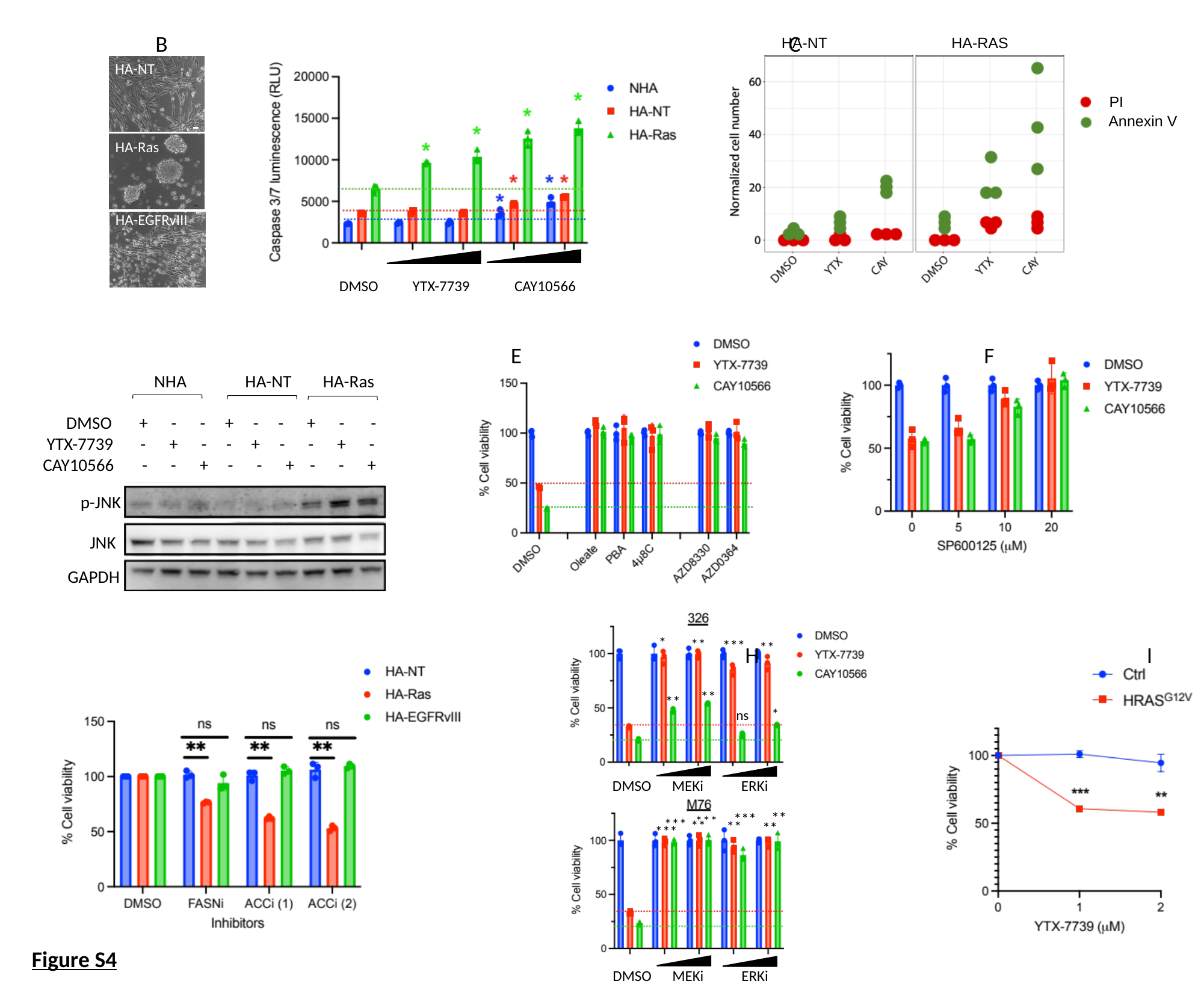

A	 B				 C
HA-NT
HA-RAS
HA-NT
HA-Ras
HA-EGFRvIII
PI
Annexin V
DMSO YTX-7739 CAY10566
D		 						 E			 	 F
NHA HA-NT HA-Ras
 DMSO + - - + - - + - -
 YTX-7739 - + - - + - - + -
 CAY10566 - - + - - + - - +
p-JNK
JNK
GAPDH
DMSO MEKi ERKi
DMSO MEKi ERKi
*
**
***
**
**
**
*
ns
**
***
***
***
**
**
**
***
G										 H I
Figure S4

### Slide 9
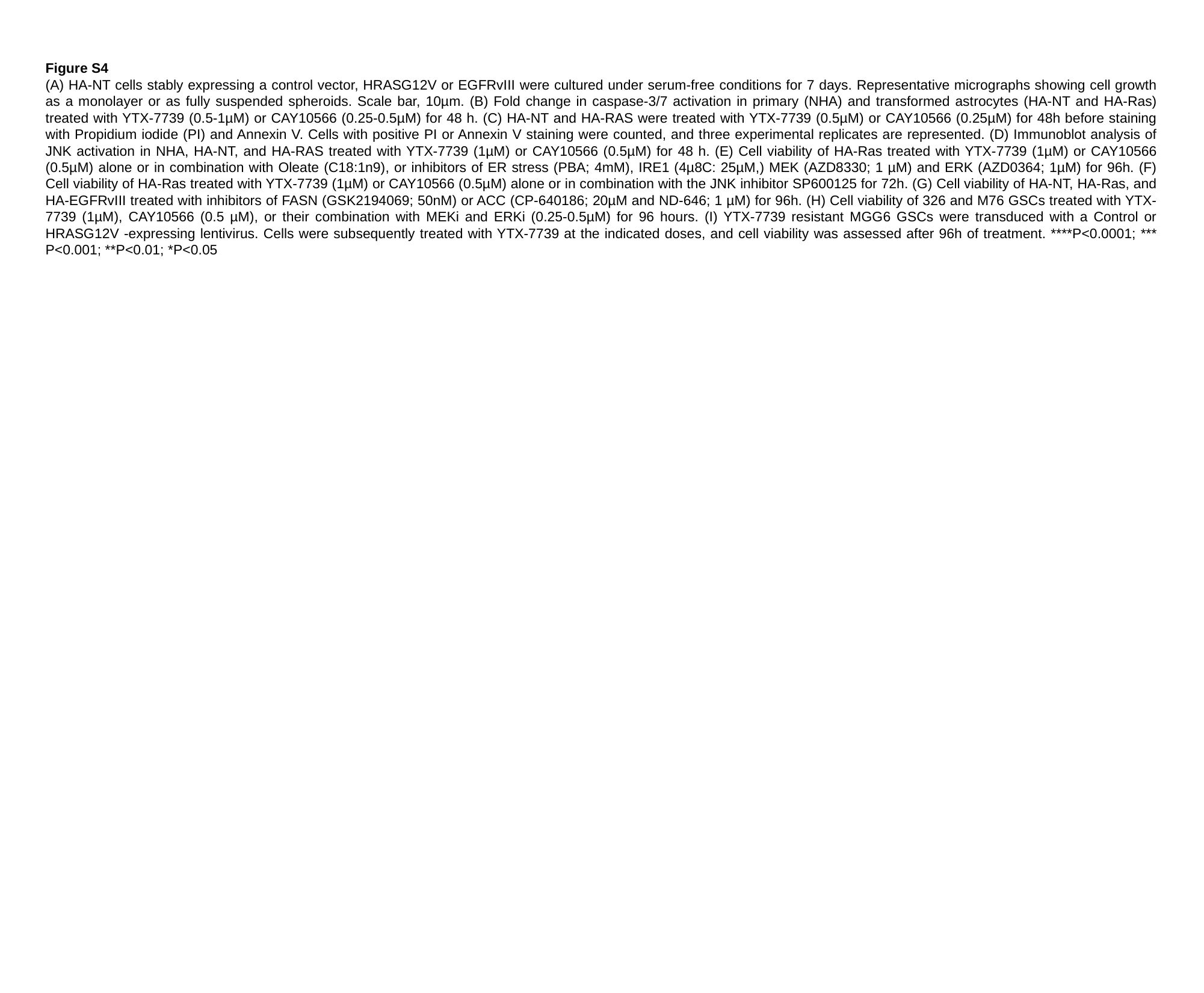

Figure S4
(A) HA-NT cells stably expressing a control vector, HRASG12V or EGFRvIII were cultured under serum-free conditions for 7 days. Representative micrographs showing cell growth as a monolayer or as fully suspended spheroids. Scale bar, 10µm. (B) Fold change in caspase-3/7 activation in primary (NHA) and transformed astrocytes (HA-NT and HA-Ras) treated with YTX-7739 (0.5-1µM) or CAY10566 (0.25-0.5µM) for 48 h. (C) HA-NT and HA-RAS were treated with YTX-7739 (0.5µM) or CAY10566 (0.25µM) for 48h before staining with Propidium iodide (PI) and Annexin V. Cells with positive PI or Annexin V staining were counted, and three experimental replicates are represented. (D) Immunoblot analysis of JNK activation in NHA, HA-NT, and HA-RAS treated with YTX-7739 (1µM) or CAY10566 (0.5µM) for 48 h. (E) Cell viability of HA-Ras treated with YTX-7739 (1µM) or CAY10566 (0.5µM) alone or in combination with Oleate (C18:1n9), or inhibitors of ER stress (PBA; 4mM), IRE1 (4µ8C: 25µM,) MEK (AZD8330; 1 µM) and ERK (AZD0364; 1µM) for 96h. (F) Cell viability of HA-Ras treated with YTX-7739 (1µM) or CAY10566 (0.5µM) alone or in combination with the JNK inhibitor SP600125 for 72h. (G) Cell viability of HA-NT, HA-Ras, and HA-EGFRvIII treated with inhibitors of FASN (GSK2194069; 50nM) or ACC (CP-640186; 20µM and ND-646; 1 µM) for 96h. (H) Cell viability of 326 and M76 GSCs treated with YTX-7739 (1µM), CAY10566 (0.5 µM), or their combination with MEKi and ERKi (0.25-0.5µM) for 96 hours. (I) YTX-7739 resistant MGG6 GSCs were transduced with a Control or HRASG12V -expressing lentivirus. Cells were subsequently treated with YTX-7739 at the indicated doses, and cell viability was assessed after 96h of treatment. ****P<0.0001; *** P<0.001; **P<0.01; *P<0.05

### Slide 10
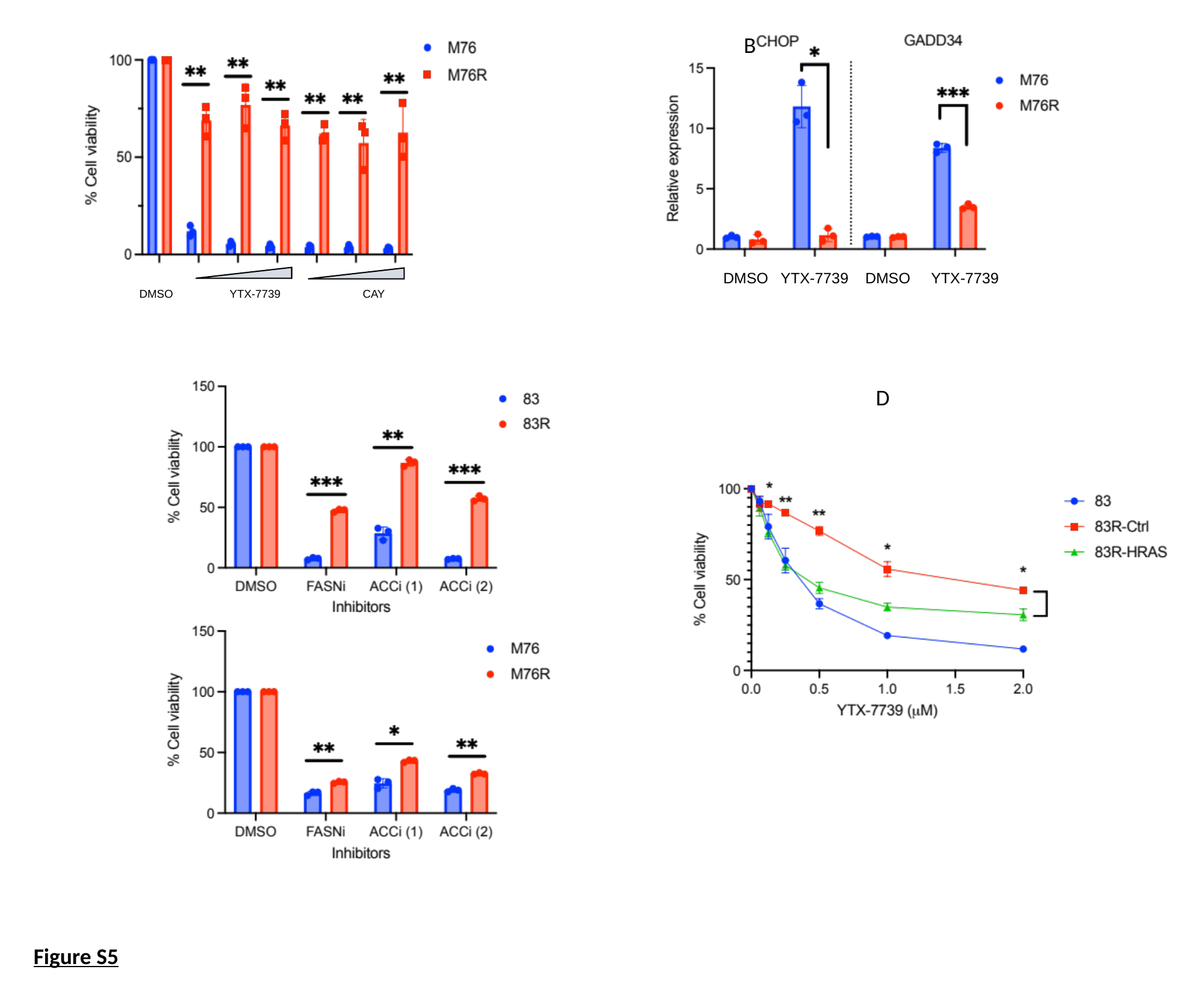

A			 				 B
DMSO YTX-7739 CAY
DMSO YTX-7739 DMSO YTX-7739
C														D
Figure S5

### Slide 11
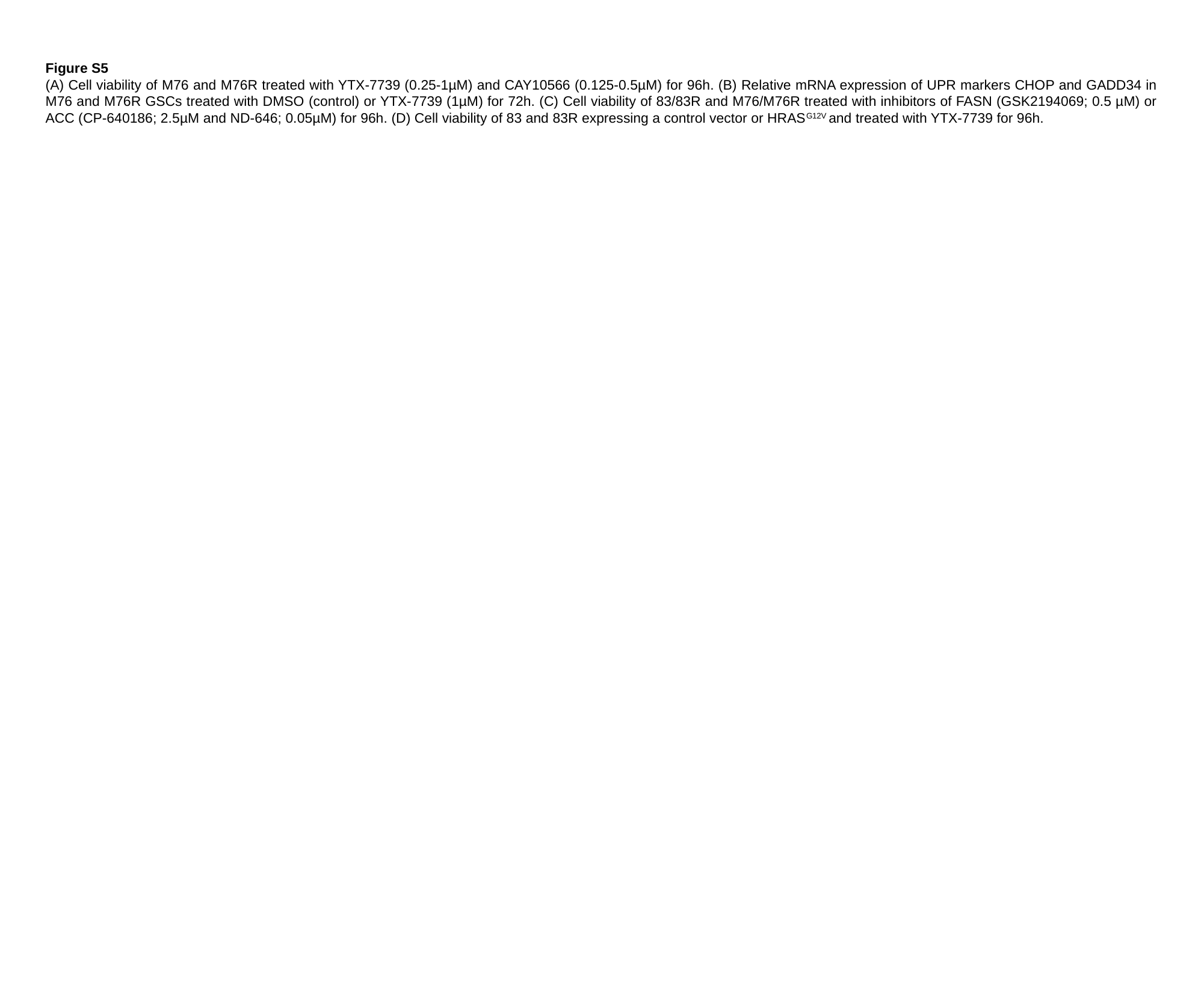

Figure S5
(A) Cell viability of M76 and M76R treated with YTX-7739 (0.25-1µM) and CAY10566 (0.125-0.5µM) for 96h. (B) Relative mRNA expression of UPR markers CHOP and GADD34 in M76 and M76R GSCs treated with DMSO (control) or YTX-7739 (1µM) for 72h. (C) Cell viability of 83/83R and M76/M76R treated with inhibitors of FASN (GSK2194069; 0.5 µM) or ACC (CP-640186; 2.5µM and ND-646; 0.05µM) for 96h. (D) Cell viability of 83 and 83R expressing a control vector or HRAS^G12V^ and treated with YTX-7739 for 96h.

### Slide 12
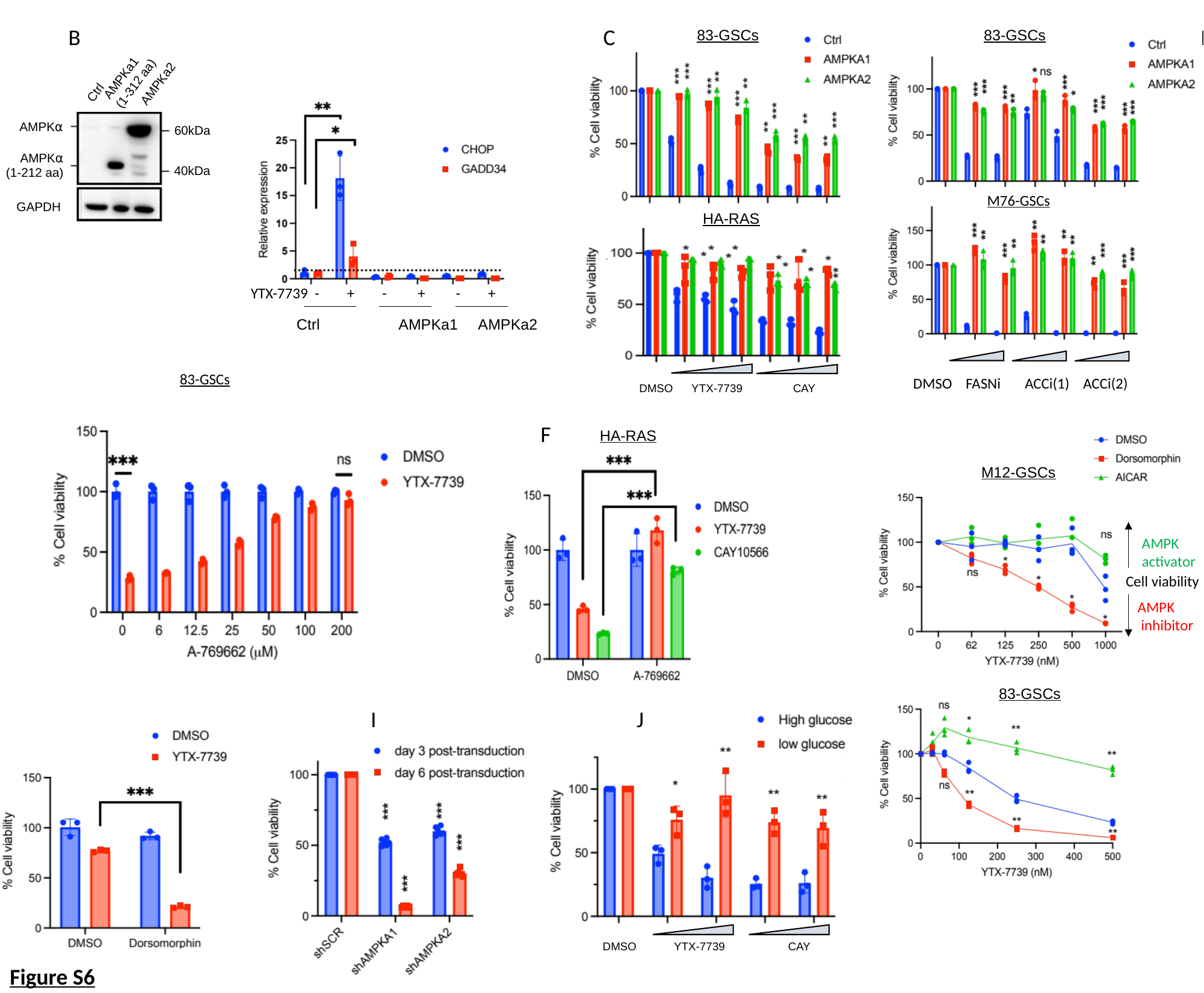

A			 B	 					C						 D
83-GSCs
HA-RAS
DMSO YTX-7739 CAY
83-GSCs
M76-GSCs
DMSO FASNi ACCi(1) ACCi(2)
AMPKa1
(1-312 aa)
AMPKa2
Ctrl
AMPK⍺
60kDa
AMPK⍺
(1-212 aa)
40kDa
GAPDH
 YTX-7739 - + - + - +
Ctrl	 AMPKa1 AMPKa2
83-GSCs
E										 F								G
HA-RAS
M12-GSCs
AMPK
activator
Cell viability
AMPK
 inhibitor
83-GSCs
H		 			 I J
DMSO YTX-7739 CAY
Figure S6

### Slide 13
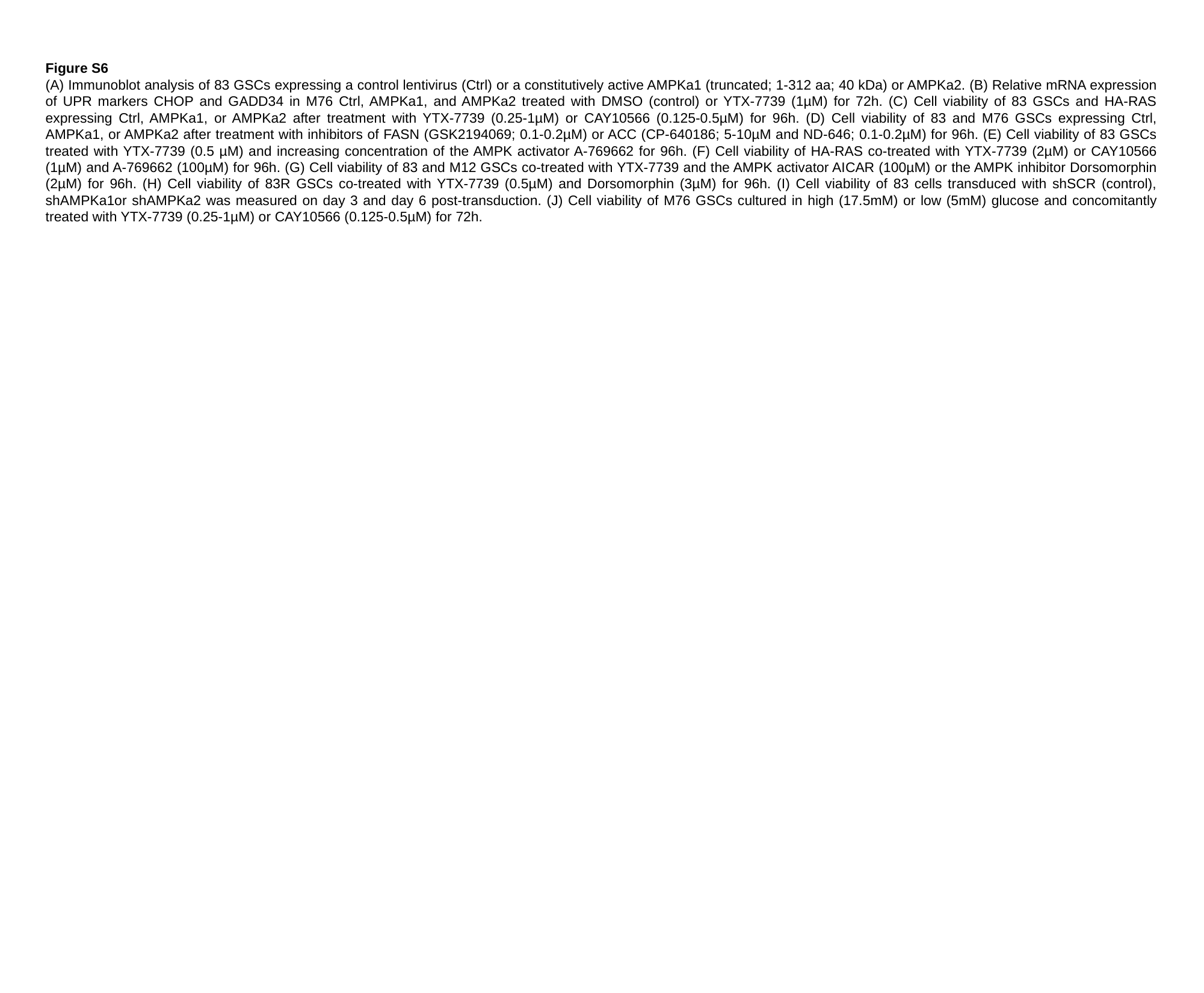

Figure S6
(A) Immunoblot analysis of 83 GSCs expressing a control lentivirus (Ctrl) or a constitutively active AMPKa1 (truncated; 1-312 aa; 40 kDa) or AMPKa2. (B) Relative mRNA expression of UPR markers CHOP and GADD34 in M76 Ctrl, AMPKa1, and AMPKa2 treated with DMSO (control) or YTX-7739 (1µM) for 72h. (C) Cell viability of 83 GSCs and HA-RAS expressing Ctrl, AMPKa1, or AMPKa2 after treatment with YTX-7739 (0.25-1µM) or CAY10566 (0.125-0.5µM) for 96h. (D) Cell viability of 83 and M76 GSCs expressing Ctrl, AMPKa1, or AMPKa2 after treatment with inhibitors of FASN (GSK2194069; 0.1-0.2µM) or ACC (CP-640186; 5-10µM and ND-646; 0.1-0.2µM) for 96h. (E) Cell viability of 83 GSCs treated with YTX-7739 (0.5 µM) and increasing concentration of the AMPK activator A-769662 for 96h. (F) Cell viability of HA-RAS co-treated with YTX-7739 (2µM) or CAY10566 (1µM) and A-769662 (100µM) for 96h. (G) Cell viability of 83 and M12 GSCs co-treated with YTX-7739 and the AMPK activator AICAR (100µM) or the AMPK inhibitor Dorsomorphin (2µM) for 96h. (H) Cell viability of 83R GSCs co-treated with YTX-7739 (0.5µM) and Dorsomorphin (3µM) for 96h. (I) Cell viability of 83 cells transduced with shSCR (control), shAMPKa1or shAMPKa2 was measured on day 3 and day 6 post-transduction. (J) Cell viability of M76 GSCs cultured in high (17.5mM) or low (5mM) glucose and concomitantly treated with YTX-7739 (0.25-1µM) or CAY10566 (0.125-0.5µM) for 72h.

### Slide 14
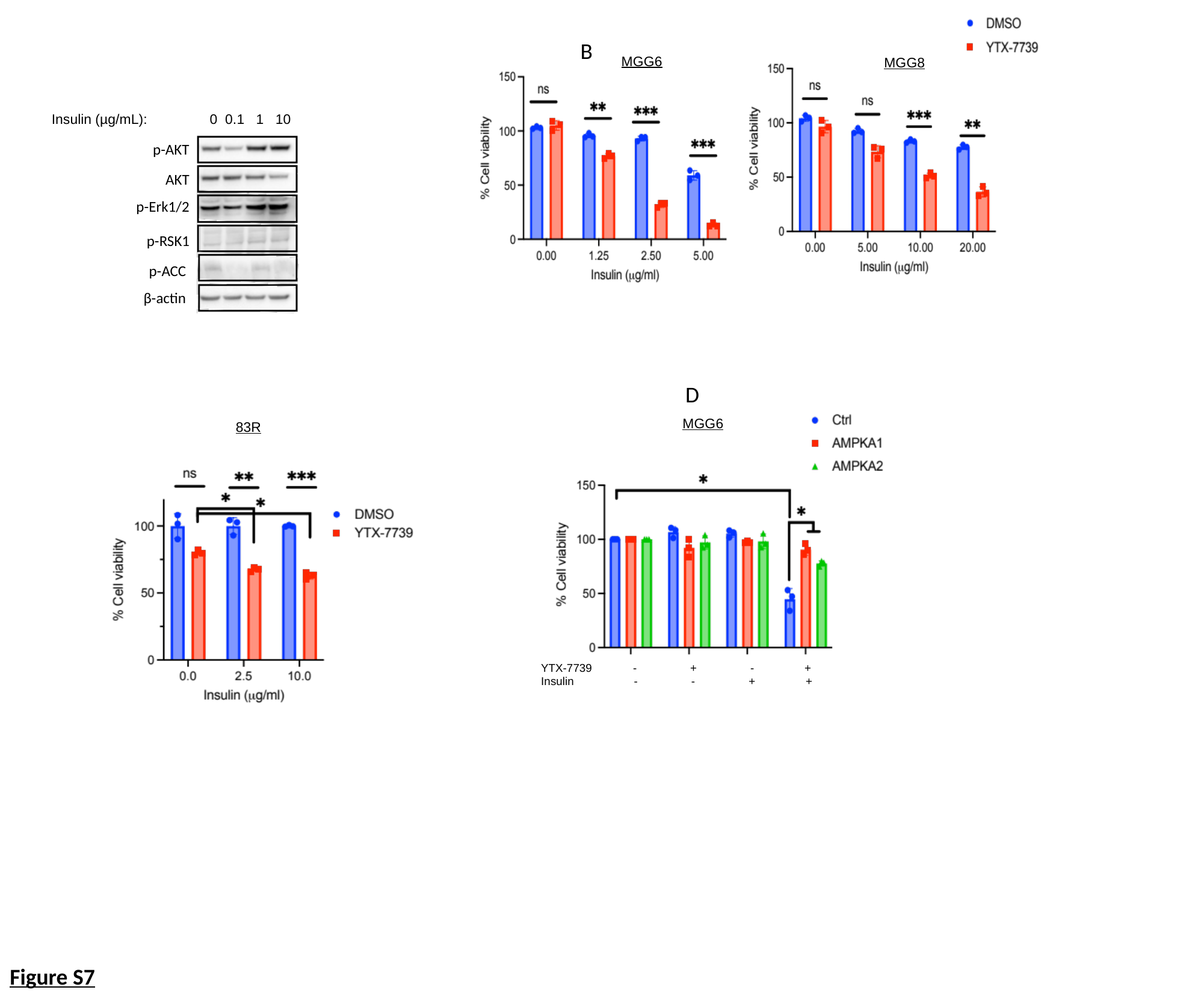

MGG6
MGG8
A									B
Insulin (µg/mL): 0 0.1 1 10
p-AKT
AKT
p-Erk1/2
p-RSK1
p-ACC
β-actin
C									 D
YTX-7739 - + - +
Insulin - - + +
MGG6
83R
Figure S7

### Slide 15
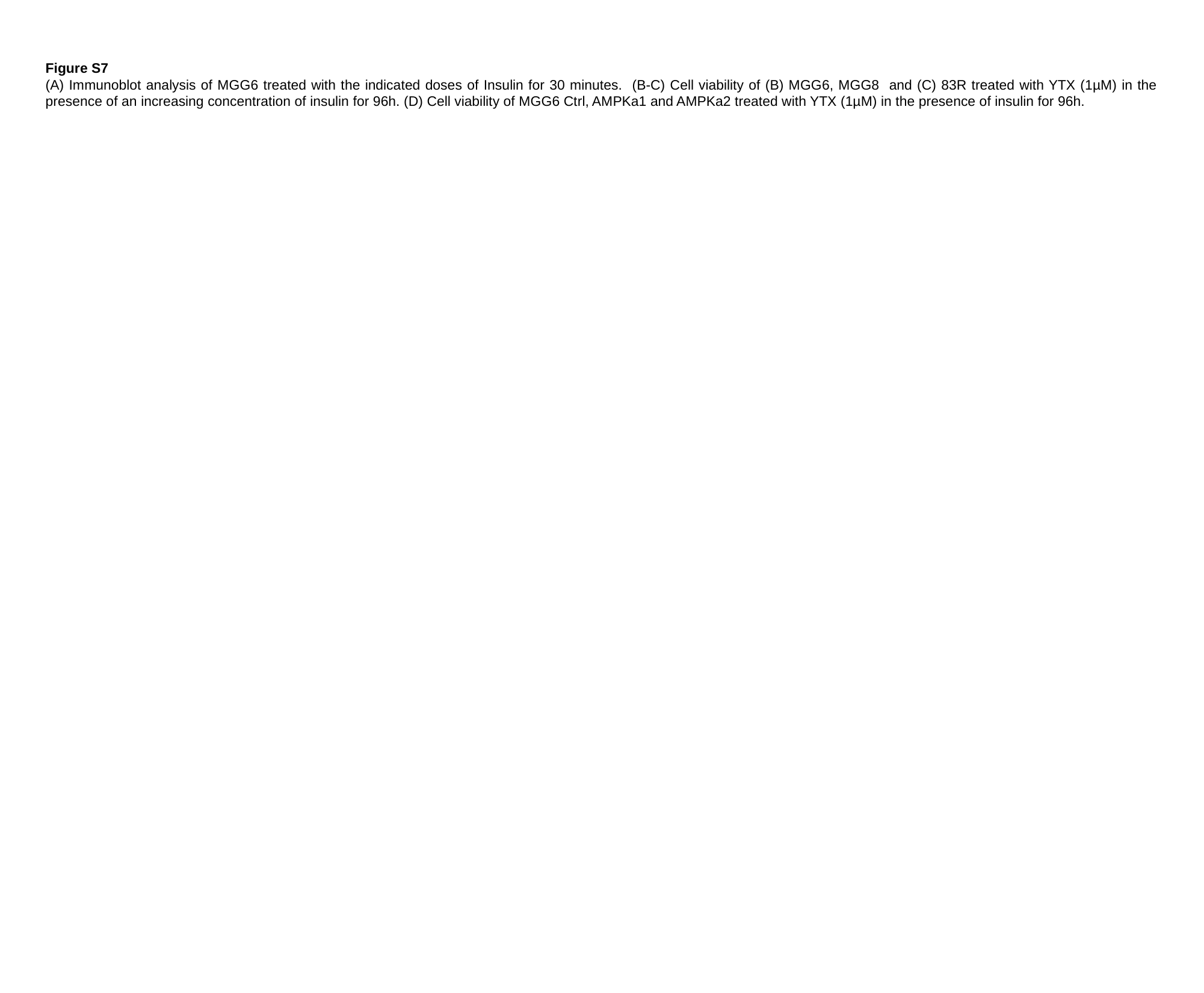

Figure S7
(A) Immunoblot analysis of MGG6 treated with the indicated doses of Insulin for 30 minutes. (B-C) Cell viability of (B) MGG6, MGG8 and (C) 83R treated with YTX (1µM) in the presence of an increasing concentration of insulin for 96h. (D) Cell viability of MGG6 Ctrl, AMPKa1 and AMPKa2 treated with YTX (1µM) in the presence of insulin for 96h.

### Slide 16
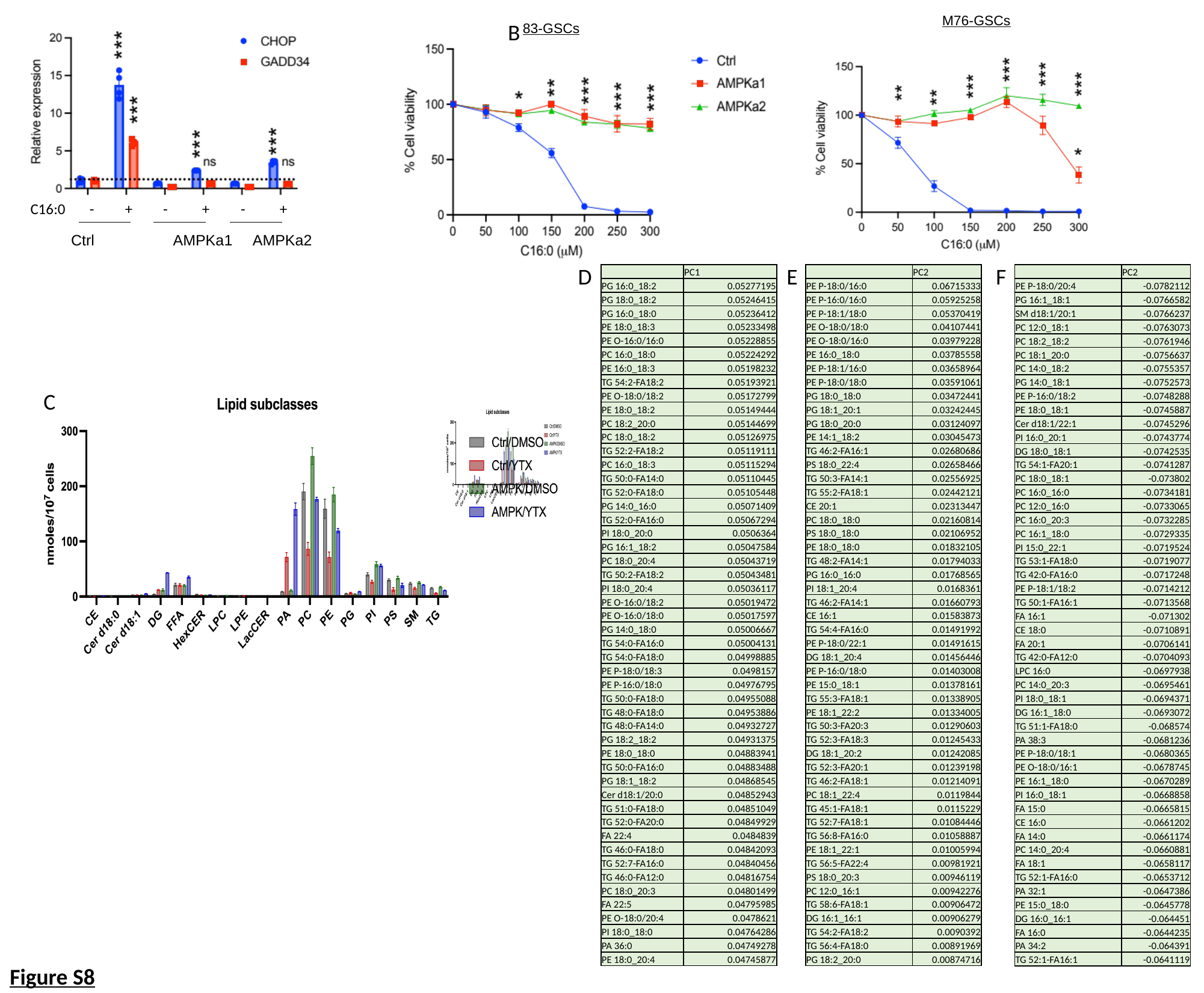

C16:0 - + - + - +
Ctrl	 AMPKa1 AMPKa2
M76-GSCs
83-GSCs
A							 B
F
D
E
| | PC1 |
| --- | --- |
| PG 16:0\_18:2 | 0.05277195 |
| PG 18:0\_18:2 | 0.05246415 |
| PG 16:0\_18:0 | 0.05236412 |
| PE 18:0\_18:3 | 0.05233498 |
| PE O-16:0/16:0 | 0.05228855 |
| PC 16:0\_18:0 | 0.05224292 |
| PE 16:0\_18:3 | 0.05198232 |
| TG 54:2-FA18:2 | 0.05193921 |
| PE O-18:0/18:2 | 0.05172799 |
| PE 18:0\_18:2 | 0.05149444 |
| PC 18:2\_20:0 | 0.05144699 |
| PC 18:0\_18:2 | 0.05126975 |
| TG 52:2-FA18:2 | 0.05119111 |
| PC 16:0\_18:3 | 0.05115294 |
| TG 50:0-FA14:0 | 0.05110445 |
| TG 52:0-FA18:0 | 0.05105448 |
| PG 14:0\_16:0 | 0.05071409 |
| TG 52:0-FA16:0 | 0.05067294 |
| PI 18:0\_20:0 | 0.0506364 |
| PG 16:1\_18:2 | 0.05047584 |
| PC 18:0\_20:4 | 0.05043719 |
| TG 50:2-FA18:2 | 0.05043481 |
| PI 18:0\_20:4 | 0.05036117 |
| PE O-16:0/18:2 | 0.05019472 |
| PE O-16:0/18:0 | 0.05017597 |
| PG 14:0\_18:0 | 0.05006667 |
| TG 54:0-FA16:0 | 0.05004131 |
| TG 54:0-FA18:0 | 0.04998885 |
| PE P-18:0/18:3 | 0.0498157 |
| PE P-16:0/18:0 | 0.04976795 |
| TG 50:0-FA18:0 | 0.04955088 |
| TG 48:0-FA18:0 | 0.04953886 |
| TG 48:0-FA14:0 | 0.04932727 |
| PG 18:2\_18:2 | 0.04931375 |
| PE 18:0\_18:0 | 0.04883941 |
| TG 50:0-FA16:0 | 0.04883488 |
| PG 18:1\_18:2 | 0.04868545 |
| Cer d18:1/20:0 | 0.04852943 |
| TG 51:0-FA18:0 | 0.04851049 |
| TG 52:0-FA20:0 | 0.04849929 |
| FA 22:4 | 0.0484839 |
| TG 46:0-FA18:0 | 0.04842093 |
| TG 52:7-FA16:0 | 0.04840456 |
| TG 46:0-FA12:0 | 0.04816754 |
| PC 18:0\_20:3 | 0.04801499 |
| FA 22:5 | 0.04795985 |
| PE O-18:0/20:4 | 0.0478621 |
| PI 18:0\_18:0 | 0.04764286 |
| PA 36:0 | 0.04749278 |
| PE 18:0\_20:4 | 0.04745877 |
| | PC2 |
| --- | --- |
| PE P-18:0/16:0 | 0.06715333 |
| PE P-16:0/16:0 | 0.05925258 |
| PE P-18:1/18:0 | 0.05370419 |
| PE O-18:0/18:0 | 0.04107441 |
| PE O-18:0/16:0 | 0.03979228 |
| PE 16:0\_18:0 | 0.03785558 |
| PE P-18:1/16:0 | 0.03658964 |
| PE P-18:0/18:0 | 0.03591061 |
| PG 18:0\_18:0 | 0.03472441 |
| PG 18:1\_20:1 | 0.03242445 |
| PG 18:0\_20:0 | 0.03124097 |
| PE 14:1\_18:2 | 0.03045473 |
| TG 46:2-FA16:1 | 0.02680686 |
| PS 18:0\_22:4 | 0.02658466 |
| TG 50:3-FA14:1 | 0.02556925 |
| TG 55:2-FA18:1 | 0.02442121 |
| CE 20:1 | 0.02313447 |
| PC 18:0\_18:0 | 0.02160814 |
| PS 18:0\_18:0 | 0.02106952 |
| PE 18:0\_18:0 | 0.01832105 |
| TG 48:2-FA14:1 | 0.01794033 |
| PG 16:0\_16:0 | 0.01768565 |
| PI 18:1\_20:4 | 0.0168361 |
| TG 46:2-FA14:1 | 0.01660793 |
| CE 16:1 | 0.01583873 |
| TG 54:4-FA16:0 | 0.01491992 |
| PE P-18:0/22:1 | 0.01491615 |
| DG 18:1\_20:4 | 0.01456446 |
| PE P-16:0/18:0 | 0.01403008 |
| PE 15:0\_18:1 | 0.01378161 |
| TG 55:3-FA18:1 | 0.01338905 |
| PE 18:1\_22:2 | 0.01334005 |
| TG 50:3-FA20:3 | 0.01290603 |
| TG 52:3-FA18:3 | 0.01245433 |
| DG 18:1\_20:2 | 0.01242085 |
| TG 52:3-FA20:1 | 0.01239198 |
| TG 46:2-FA18:1 | 0.01214091 |
| PC 18:1\_22:4 | 0.0119844 |
| TG 45:1-FA18:1 | 0.0115229 |
| TG 52:7-FA18:1 | 0.01084446 |
| TG 56:8-FA16:0 | 0.01058887 |
| PE 18:1\_22:1 | 0.01005994 |
| TG 56:5-FA22:4 | 0.00981921 |
| PS 18:0\_20:3 | 0.00946119 |
| PC 12:0\_16:1 | 0.00942276 |
| TG 58:6-FA18:1 | 0.00906472 |
| DG 16:1\_16:1 | 0.00906279 |
| TG 54:2-FA18:2 | 0.0090392 |
| TG 56:4-FA18:0 | 0.00891969 |
| PG 18:2\_20:0 | 0.00874716 |
| | PC2 |
| --- | --- |
| PE P-18:0/20:4 | -0.0782112 |
| PG 16:1\_18:1 | -0.0766582 |
| SM d18:1/20:1 | -0.0766237 |
| PC 12:0\_18:1 | -0.0763073 |
| PC 18:2\_18:2 | -0.0761946 |
| PC 18:1\_20:0 | -0.0756637 |
| PC 14:0\_18:2 | -0.0755357 |
| PG 14:0\_18:1 | -0.0752573 |
| PE P-16:0/18:2 | -0.0748288 |
| PE 18:0\_18:1 | -0.0745887 |
| Cer d18:1/22:1 | -0.0745296 |
| PI 16:0\_20:1 | -0.0743774 |
| DG 18:0\_18:1 | -0.0742535 |
| TG 54:1-FA20:1 | -0.0741287 |
| PC 18:0\_18:1 | -0.073802 |
| PC 16:0\_16:0 | -0.0734181 |
| PC 12:0\_16:0 | -0.0733065 |
| PC 16:0\_20:3 | -0.0732285 |
| PC 16:1\_18:0 | -0.0729335 |
| PI 15:0\_22:1 | -0.0719524 |
| TG 53:1-FA18:0 | -0.0719077 |
| TG 42:0-FA16:0 | -0.0717248 |
| PE P-18:1/18:2 | -0.0714212 |
| TG 50:1-FA16:1 | -0.0713568 |
| FA 16:1 | -0.071302 |
| CE 18:0 | -0.0710891 |
| FA 20:1 | -0.0706141 |
| TG 42:0-FA12:0 | -0.0704093 |
| LPC 16:0 | -0.0697938 |
| PC 14:0\_20:3 | -0.0695461 |
| PI 18:0\_18:1 | -0.0694371 |
| DG 16:1\_18:0 | -0.0693072 |
| TG 51:1-FA18:0 | -0.068574 |
| PA 38:3 | -0.0681236 |
| PE P-18:0/18:1 | -0.0680365 |
| PE O-18:0/16:1 | -0.0678745 |
| PE 16:1\_18:0 | -0.0670289 |
| PI 16:0\_18:1 | -0.0668858 |
| FA 15:0 | -0.0665815 |
| CE 16:0 | -0.0661202 |
| FA 14:0 | -0.0661174 |
| PC 14:0\_20:4 | -0.0660881 |
| FA 18:1 | -0.0658117 |
| TG 52:1-FA16:0 | -0.0653712 |
| PA 32:1 | -0.0647386 |
| PE 15:0\_18:0 | -0.0645778 |
| DG 16:0\_16:1 | -0.064451 |
| FA 16:0 | -0.0644235 |
| PA 34:2 | -0.064391 |
| TG 52:1-FA16:1 | -0.0641119 |
E
C
Figure S8

### Slide 17
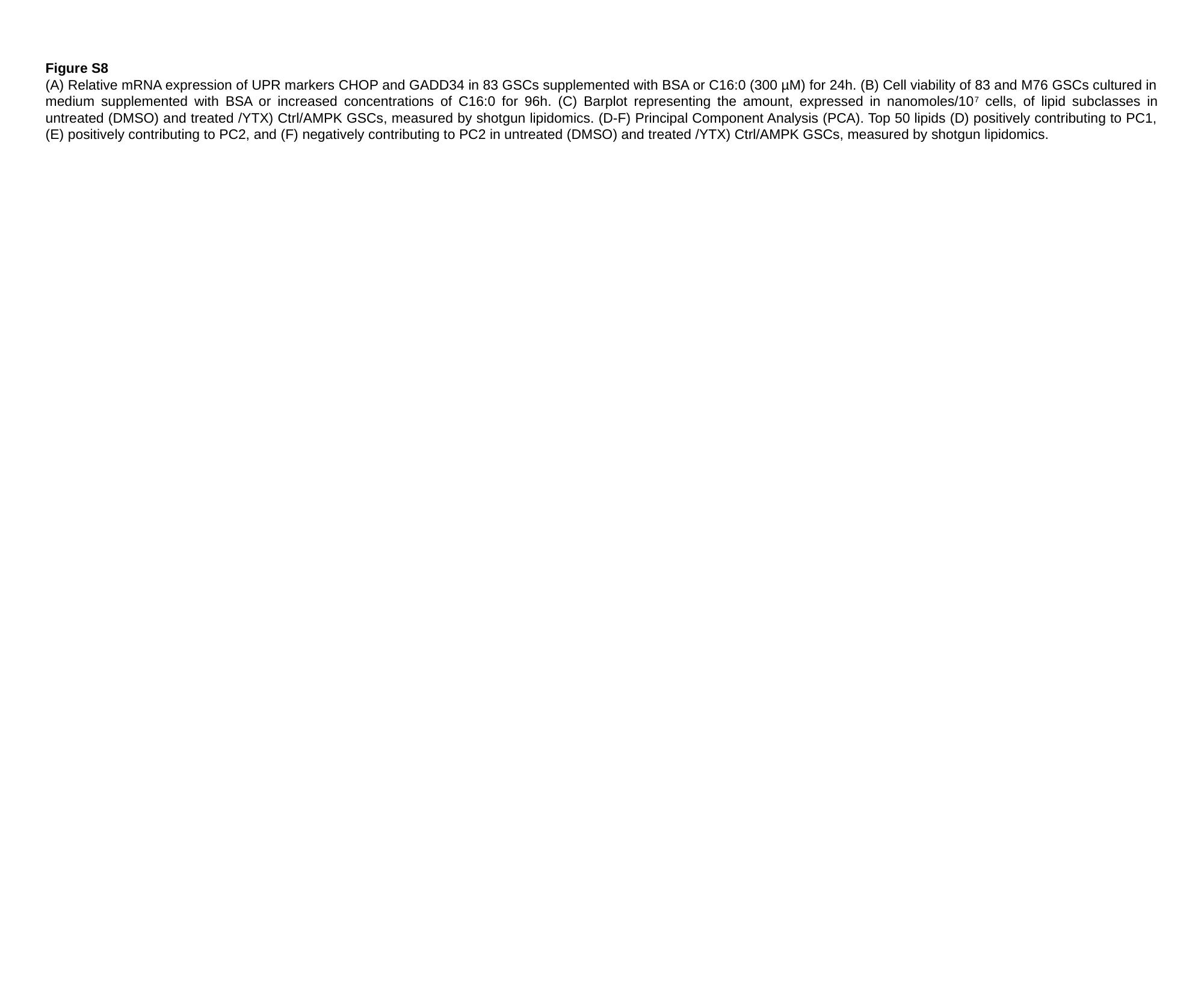

Figure S8
(A) Relative mRNA expression of UPR markers CHOP and GADD34 in 83 GSCs supplemented with BSA or C16:0 (300 µM) for 24h. (B) Cell viability of 83 and M76 GSCs cultured in medium supplemented with BSA or increased concentrations of C16:0 for 96h. (C) Barplot representing the amount, expressed in nanomoles/10^7^ cells, of lipid subclasses in untreated (DMSO) and treated /YTX) Ctrl/AMPK GSCs, measured by shotgun lipidomics. (D-F) Principal Component Analysis (PCA). Top 50 lipids (D) positively contributing to PC1, (E) positively contributing to PC2, and (F) negatively contributing to PC2 in untreated (DMSO) and treated /YTX) Ctrl/AMPK GSCs, measured by shotgun lipidomics.

### Slide 18
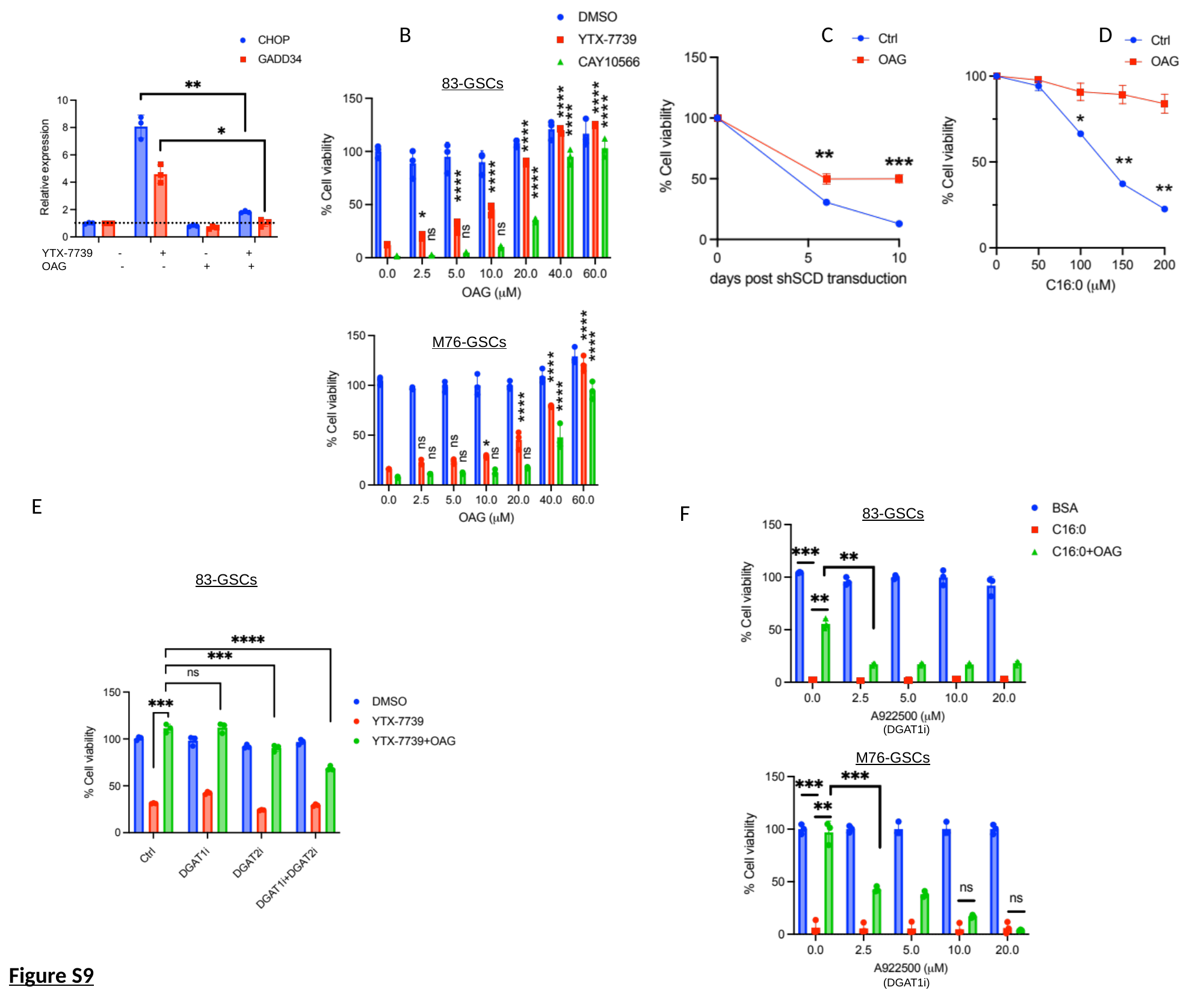

A					 B	 		 C D
83-GSCs
M76-GSCs
YTX-7739 - + - +
OAG - - + +
E
F
83-GSCs
83-GSCs
(DGAT1i)
M76-GSCs
Figure S9
(DGAT1i)

### Slide 19
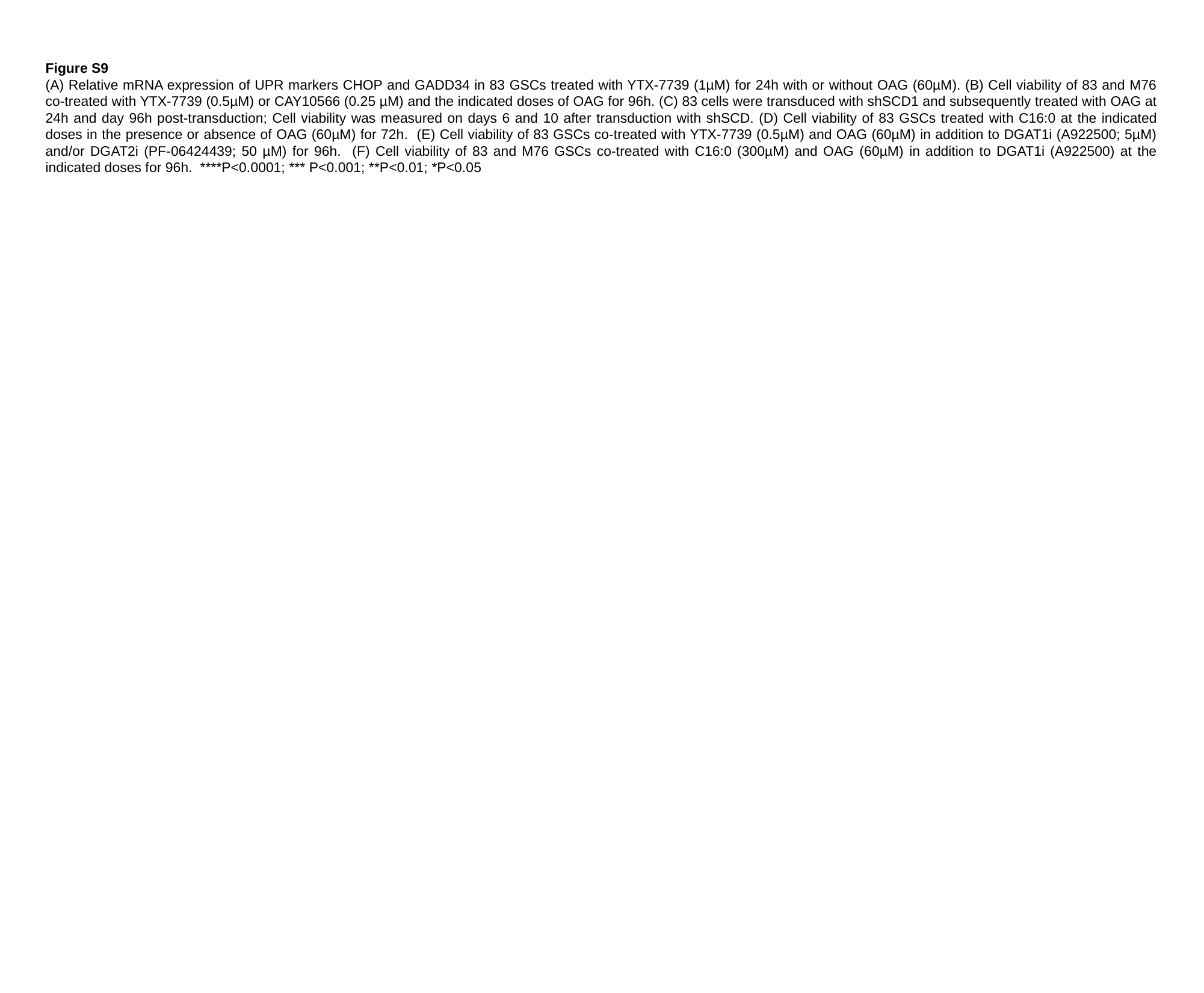

Figure S9
(A) Relative mRNA expression of UPR markers CHOP and GADD34 in 83 GSCs treated with YTX-7739 (1µM) for 24h with or without OAG (60µM). (B) Cell viability of 83 and M76 co-treated with YTX-7739 (0.5µM) or CAY10566 (0.25 µM) and the indicated doses of OAG for 96h. (C) 83 cells were transduced with shSCD1 and subsequently treated with OAG at 24h and day 96h post-transduction; Cell viability was measured on days 6 and 10 after transduction with shSCD. (D) Cell viability of 83 GSCs treated with C16:0 at the indicated doses in the presence or absence of OAG (60µM) for 72h. (E) Cell viability of 83 GSCs co-treated with YTX-7739 (0.5µM) and OAG (60µM) in addition to DGAT1i (A922500; 5µM) and/or DGAT2i (PF-06424439; 50 µM) for 96h. (F) Cell viability of 83 and M76 GSCs co-treated with C16:0 (300µM) and OAG (60µM) in addition to DGAT1i (A922500) at the indicated doses for 96h. ****P<0.0001; *** P<0.001; **P<0.01; *P<0.05
